# Assessing the impact of data uncertainty on remote sensing estimation of plant functional diversity

**DOI:** 10.64898/2026.09.21.753123

**Authors:** Javier Pacheco Labrador, Christian Rossi, Maria J. Santos

## Abstract

Remote sensing mapping of plant functional diversity (PFD) is increasingly used at the science-policy interface. Yet assessments of uncertainty and quality standards for these products are seldom provided, as the impact of measurement uncertainty on PFD estimates, i.e., the variability across pixels, is still poorly understood. We use trait maps and hyperspectral imagery simulations from the Biodiversity Observing System Simulation Experiment (BOSSE) to assess the impact of different types of uncertainty on the estimation PFD. Results show that uncertainties stemming from random errors affect the estimation of PFD (up to three times) more than those arising from systematic errors, with spectral and spatial error correlation reducing their impact. Standardization computing PFD metrics reduces systematic errors, making in-situ-measured plant traits and remote sensing reflectance factors more robust estimators of PFD than other remote sensing proxies derived from reflectance (e.g., spectral indices or optical traits). The impact of uncertainty propagation from reflectance factors to these often-preferred secondary proxies increases with their complexity. Nonetheless, trait uncertainty can spuriously diminish differences between PFD derived from the ground (plant traits) and satellite imagery, as it can reduce uncertainty-induced or inherent biases between the PFD metrics computed from each trait (plant or spectral). This study provides new understanding of how uncertainty propagates into PFD estimates; this knowledge is relevant for optimizing the selection and defining quality requirements for PFD estimation. It also suggests that random and systematic uncertainties should be quantified separately at the reflectance factor level to guide their use and interpretation in PFD analysis.

## 1. INTRODUCTION

Plant functional diversity (PFD), defined as the spatial variability of plant traits within communities, provides a critical link between biodiversity and ecosystem functioning, making it especially relevant for monitoring and decision-making (Díaz et al., 2007). Accordingly, estimates of PFD derived from optical remote sensing data are becoming increasingly widespread (Cavender-Bares et al., 2022; Wang and Gamon, 2019). This trend is driven by advances in imaging spectroscopy and its ability to infer plant traits from reflectance signals (Cimoli et al., 2024; Ollinger, 2011; Ustin et al., 2009; Wang et al., 2019a). Estimating PFD from remote sensing data can be done from the spatial variability of remotely sensed plant trait estimates (i.e., optical traits) (Gamon et al., 2019; Ustin et al., 2009), but also from surface reflectance values and derived spectral indices (Beccari et al., 2024; Pacheco-Labrador et al., 2022; Rossi et al., 2024; Torresani et al., 2024). Therefore, recent and upcoming advances in remote sensing, particularly imaging spectroscopy missions, are expected to generalize the estimation of plant functional traits and diversity from space at global and local scales (Ma et al., 2020; Skidmore et al., 2021; Zhang et al., 2021).

As with any measurement, remotely sensed PFD estimates are inherently uncertain (Rossi et al., in press). Starting with the digital numbers recorded by the sensor’s detector and continuing through radiometric and spectral calibration, orthorectification, topographic, atmospheric, and anisotropy correction, multiple sources of uncertainty can be introduced at each step of the processing chain (Bachmann et al., 2015; Gorroño et al., 2024; Graf et al., 2023; Hueni et al., 2017; K. Thome, 2016; Martínez-Ferrer et al., 2022). Methodological choices or limitations can also increase uncertainty (Torresani et al., 2024; Wallis et al., 2025). Yet, the propagation of these uncertainties into PFD estimates remains rarely implemented, explicitly considered, or meaningfully interpreted, limiting our ability to assess their robustness. Failing to account for uncertainties in both in-situ and remote sensing data (as well as the models that link these variables or estimate optical traits) limits evaluation of remote sensing capability to map PFD and the development and selection of robust methods and plausible diversity measurements. Beyond constraining method evaluation and selection, this omission also prevents quantifying the uncertainty in the resulting PFD estimates, making it difficult to assess their conformity to application requirements (Widlowski, 2015) with downstream repercussions for management strategies and policymaking.

The impact of different types of uncertainty on PFD estimation may differ from its impact on plant-trait estimation itself. Whereas the latter seeks accurate and precise mapping of plant traits (Wang et al., 2019a), the former, diversity estimation, focuses on the spatial variability of the functional traits (Asner et al., 2017), with multiple diversity metrics exhibiting different sensitivities to errors due to their different formulations. For example, Pacheco-Labrador et al. (2022) assessed the suitability of different diversity metrics for remote sensing data and found that uncertainties related to spatial resolution spuriously increased the correlation between in-situ plant traits and remote sensing diversity values for some metrics (e.g., functional richness). The authors also hypothesized that equifinality in the retrieval of optical traits might attenuate uncertainty in the subsequent PFD estimation.

It remains therefore unclear (a) which remote sensing proxies are more robust to uncertainty when used to estimate PFD, (b) what is the joint effect of uncertainties present in the remote sensing proxies and the in-situ plant trait measurements, and (c) to what extent the preprocessing of the plant and spectral traits can attenuate or inflate the effect of different types of uncertainty and their impacts on PFD estimation. We explore these questions using synthetic scenes that encompass plant traits and remote sensing proxies of PFD (hyperspectral reflectance, spectral indices, and optical traits) generated with the Biodiversity Observing System Simulation Experiment (BOSSE) (Pacheco-Labrador et al., 2025).

## 2. METHODS

Fig. 1 summarizes the workflow applied in this study. First, we simulated synthetic plant trait maps (i.e., radiative transfer model foliar and structural parameters) and the corresponding imaging spectroscopy reflectance using BOSSE (Pacheco-Labrador et al., 2024) (Fig. 1a). Second, we applied different types of uncertainty (*u*) (Fig. 1b) to the simulated synthetic plant trait values and spectral reflectance factors. We used the spectral reflectance factors to compute spectral vegetation indices and optical traits (reflectance-based estimates of plant traits (Feilhauer et al., 2017)), and propagated uncertainties through these calculations. Third, we computed Rao’s quadratic entropy index (*Q*_Rao_, Botta-Dukát (2005)), a frequently used functional diversity metric, for the plant traits and the remote sensing proxies (Fig. 1c). Finally, we assessed the effect of uncertainty by regressing *Q*_Rao_ computed from synthetic plant trait maps against *Q*_Rao_ computed from the different remote sensing proxies for plant traits (Fig. 1d).

**Figure 1.**
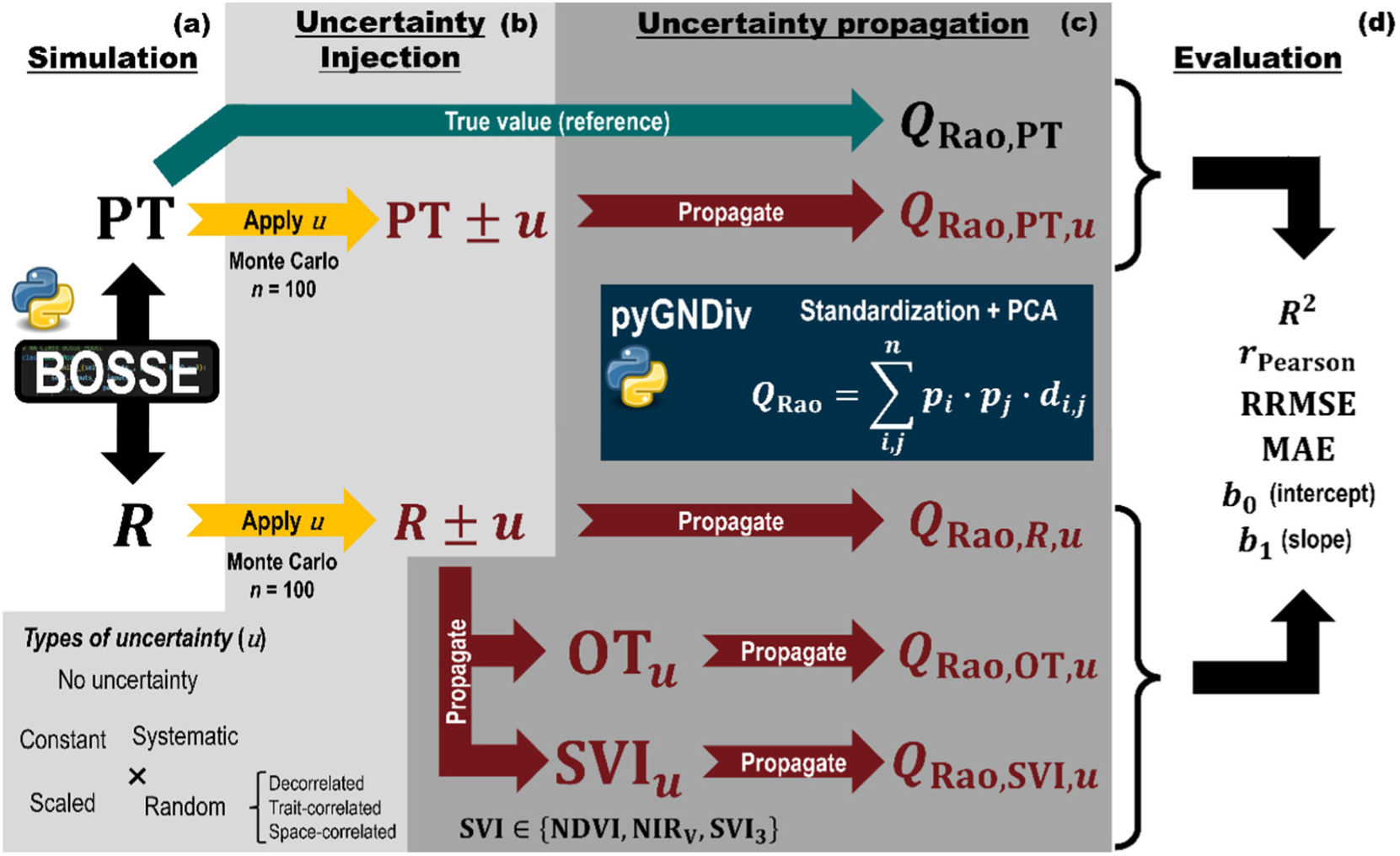
Workflow concatenating (a) the simulation of in-situ plant traits (PT) and the hyperspectral reflectance factors (*R*), (b) the addition of different types of uncertainty (*u*), (c) the propagation to secondary reflectance-based variables (i.e., optical traits (OT) and spectral vegetation indices (SVI)) and functional diversity metrics (i.e., Raos’ quadratic entropy index (*Q*_Rao_)), and (d) the comparison of functional diversity computed from plant traits and the rest of the simulated variables using coefficient of determination (*R*^2^), Pearson correlation coefficient (*r*_Pearson_), relative root mean squared error (RRMSE, [%]), mean absolute error (MAE, [-]), and the slope (*b*_1_) and bias (*b*_0_) of the relationship between predicted (*Q*_Rao_ from remote sensing estimates) - observed (*Q*_Rao_ from in-situ plant traits). The spectral indices used are the normalized difference vegetation index (NDVI), the near infrared of vegetation (NIR_v_), and a cube of three spectral indices (SVI_3_), proxies of leaf chlorophyll, carotenoids, and water content.

### 2.1 BOSSE simulations

BOSSE is a virtual scene generator capable of simulating spatially distributed species and the corresponding plant traits (Table 1), along with the corresponding physically-linked remote sensing imagery and ecosystem functions (e.g., carbon, water, and energy fluxes) by emulating the SCOPE model (van der Tol et al., 2009). BOSSE provides synthetic, comparable datasets across a wide range of situations (e.g., spatial distributions of species, taxonomic richness, inter- and intra-specific diversity, meteorological conditions, among others) that enable robust benchmarking of methods for inferring PFD from remote sensing (e.g., Pacheco-Labrador et al. (2026)). BOSSE scenes are randomly generated, but traits are constrained by covariance and bounds reported in the literature and in open-access datasets (Pacheco-Labrador et al., 2025) (Table 1). A phenological model describes the temporal evolution of plant traits as a function of meteorological conditions (Pacheco-Labrador et al., 2025).

**Table 1.**
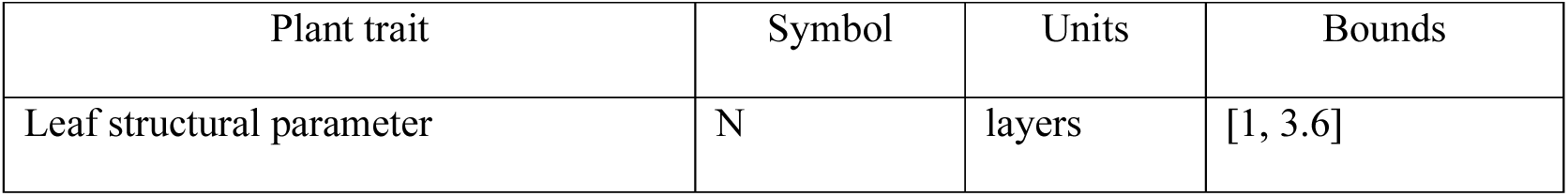
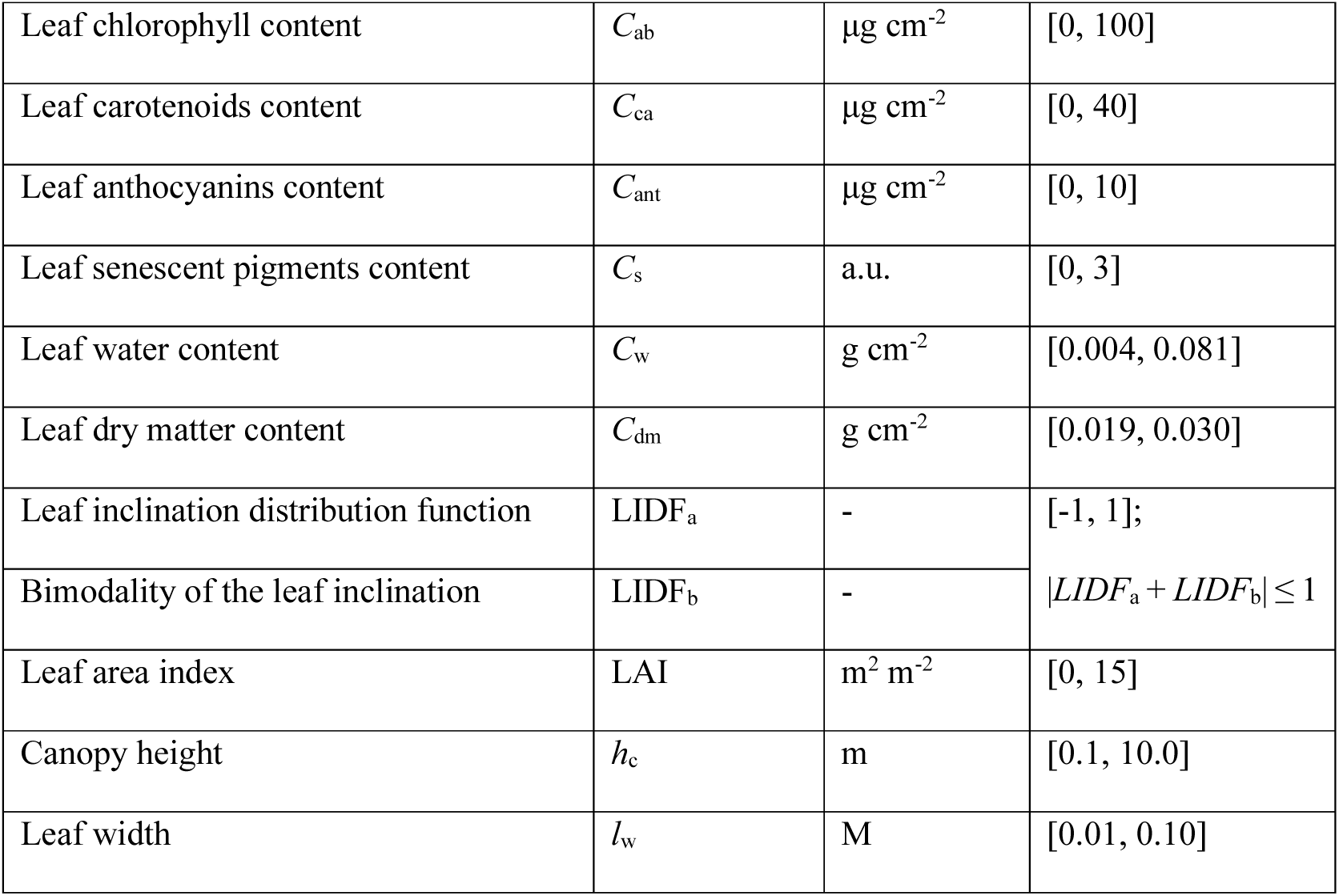
BOSSE plant (and optical) trait symbols, units, and ranges used for the simulations.

We simulated 180 synthetic scenes, each 120 by 120 pixels, using BOSSE. To do so, we used 15 different meteorological time series for each of the four climatic zones (tropical, temperate, continental, and arid), which determine the abundance of different plant functional types, and three spatial patterns (clustered, intermediate, and even) which determine their spatial distribution in the scene (Fig. 2a-c, d-f, and g-i, respectively). Each scene featured a different species richness, randomly selected from 1 to 60. For each scene, we selected four dates corresponding to the 100^th^, 85^th^, 60^th^, and 45^th^ percentiles of the scene mean phenology index (i.e., the Growing Season Index (Forkel et al., 2014), ranging from 0 to 1). Phenology affects functional diversity as it determines the fraction of bare soil pixels and the range of trait values (e.g., evergreen species alongside developing or decaying deciduous species). Hence, for each scene, we generated four reflectance maps, resulting in a total of 720 pairs of plant trait maps with corresponding remote sensing images. Plant traits and reflectance factors are non-linearly linked through radiative transfer model emulators: statistical surrogates of radiative transfer models (Gómez-Dans et al., 2016) that predict (hemispherical-directional) reflectance factors as a function of plant traits. In BOSSE, an independently trained inverse model predicts these traits from the reflectance factors (i.e., the optical traits), introducing additional model uncertainty in the retrieval.

**Figure 2.**
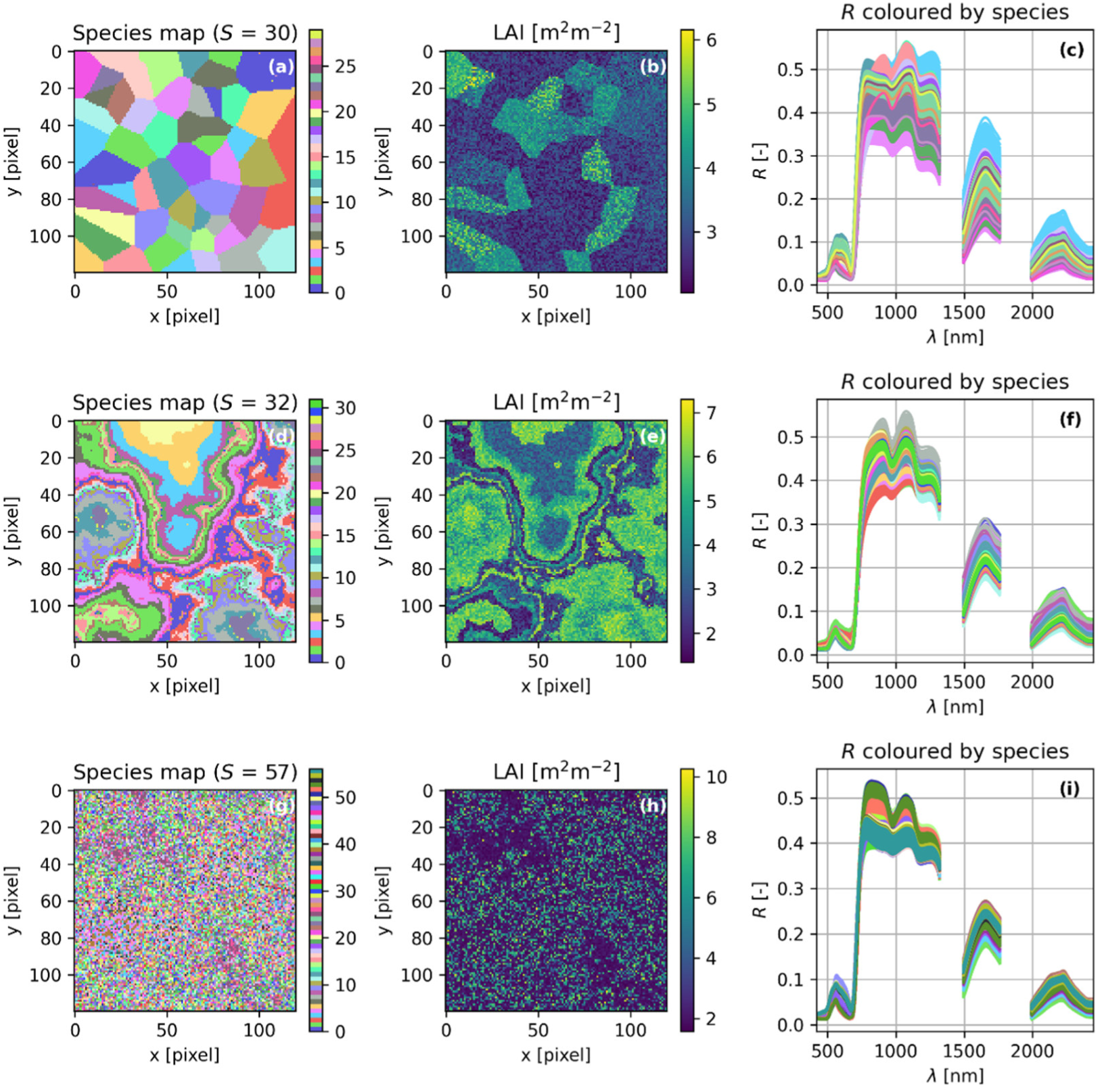
Example of three BOSSE scenes generated with clustered (first row), intermediate (middle row), and even (bottom row) spatial patterns. The figure depicts the species maps (left column), the corresponding leaf area index (middle column), and spectral reflectance factors (right column, with colors matching those of the species).

For each date and scene, we generated six synthetic sets of variables from which we separately computed functional diversity (Table 2): in-situ plant trait (PT) maps, representing an exhaustive knowledge of field plant traits, and the corresponding hyperspectral reflectance factors (*R*) featuring the EnMAP spectral configuration (i.e., 197 spectral bands after removing atmospheric water absorption regions) (Storch et al., 2023). From *R*, we first computed three sets of spectral vegetation indices (SVI): a) the normalized difference vegetation index (NDVI) (Kriegler et al., 1969), b) the near infrared of vegetation (NIR_v_) (Badgley et al., 2017), and c) a cube of three spectral indices (SVI_3_), proxies of foliar chlorophyll (*C*_ab_), carotenoids (*C*_ca_), and water (*C*_w_) content as defined in Schneider et al. (2017) (Table 2), and second, retrieved optical traits (OT, i.e., reflectance-based plant traits estimates, using a neural network integrated in BOSSE for the retrieval) (Fig. 1a-c). We simulated spectral information at midday to reflect the acquisition time of most optical sun-synchronous sensors. PT served as the in-situ reference data to compute PFD, whereas the remaining remote sensing proxies (*R*, SVIs, and OT) were used to estimate such diversity.

**Table 2.** Plant or spectral traits used to compute functional diversity.

| Trait | Symbol | Num.<br>of<br>vars. | Description |
| --- | --- | --- | --- |
| Field measurements |  |  |  |
| Plant traits | PT | 12 | Leaf structural parameters, leaf pigments (chlorophyll, carotenoids, anthocyanins, senescent), water, and dry matter content, leaf area index, leaf angle distribution parameters, leaf width, and canopy height (Table 1). They represent the “in-situ” data from which plant functional diversity is originally calculated and used as a reference for the remaining remote sensing proxies. |
| Remote sensing imagery and products |  |  |  |
| Reflectance factors | $R$ | 197 | Hemispherical-Directional Reflectance Factors featuring EnMAP spectral configuration without atmospheric water absorption bands |
| Normalized Difference Vegetation Index | NDVI | 1 | $NDVI = \frac{R_{NIR} - R_{Red}}{R_{NIR} + R_{Red}}$ |
| Near Infrared of Vegetation | $NIR_v$ | 1 | $NIR_v = R_{NIR} \cdot \frac{R_{NIR} - R_{Red}}{R_{NIR} + R_{Red}}$ |
| Vegetation indices cube (Schneider et al., 2017) | $SVI_3$ | 3 | Set of three spectral indices representing chlorophyll, carotenoids, and water content used by Schneider et al. (2017).<br>$\begin{bmatrix} C_{ab} \\ C_{ar} \\ C_w \end{bmatrix} = \begin{bmatrix} \left( \frac{1}{R_{540-560}} - \frac{1}{R_{760-800}} \right) R_{760-800} \\ \left( \frac{1}{R_{510-520}} - \frac{1}{R_{690-710}} \right) R_{760-800} \\ 1 - \frac{R_{1193}}{R_{1126}} \end{bmatrix}$ |
| Optical traits | OT | 12 | Plant traits estimated from the reflectance factors using a neural network. Presented in Table 1. |

### 2.2 Uncertainty simulation

We generated different types of data uncertainty that can occur in remote sensing and in-situ measurements (Table 3) and applied them to the plant traits and the reflectance factors. The uncertainty added to *R* was then propagated to the SVIs and the retrieval of OT (Fig. 1b,c). Following Vasquez and Whiting (2005), we simulated uncertainties stemming from systematic errors (*e*_sys_, bias) and random errors (*e*_rnd_, noise) through two probability distribution functions (PDFs): Uniform for systematic errors and Gaussian for random errors. Importantly, error (*e*) refers to a single realization of deviation from the true value, whereas uncertainty (*u*) describes the distribution of errors. We used a Monte Carlo approach, drawing 100 realizations of possible error values, and added the values to generate perturbed PT and *R*.

**Table 3.**
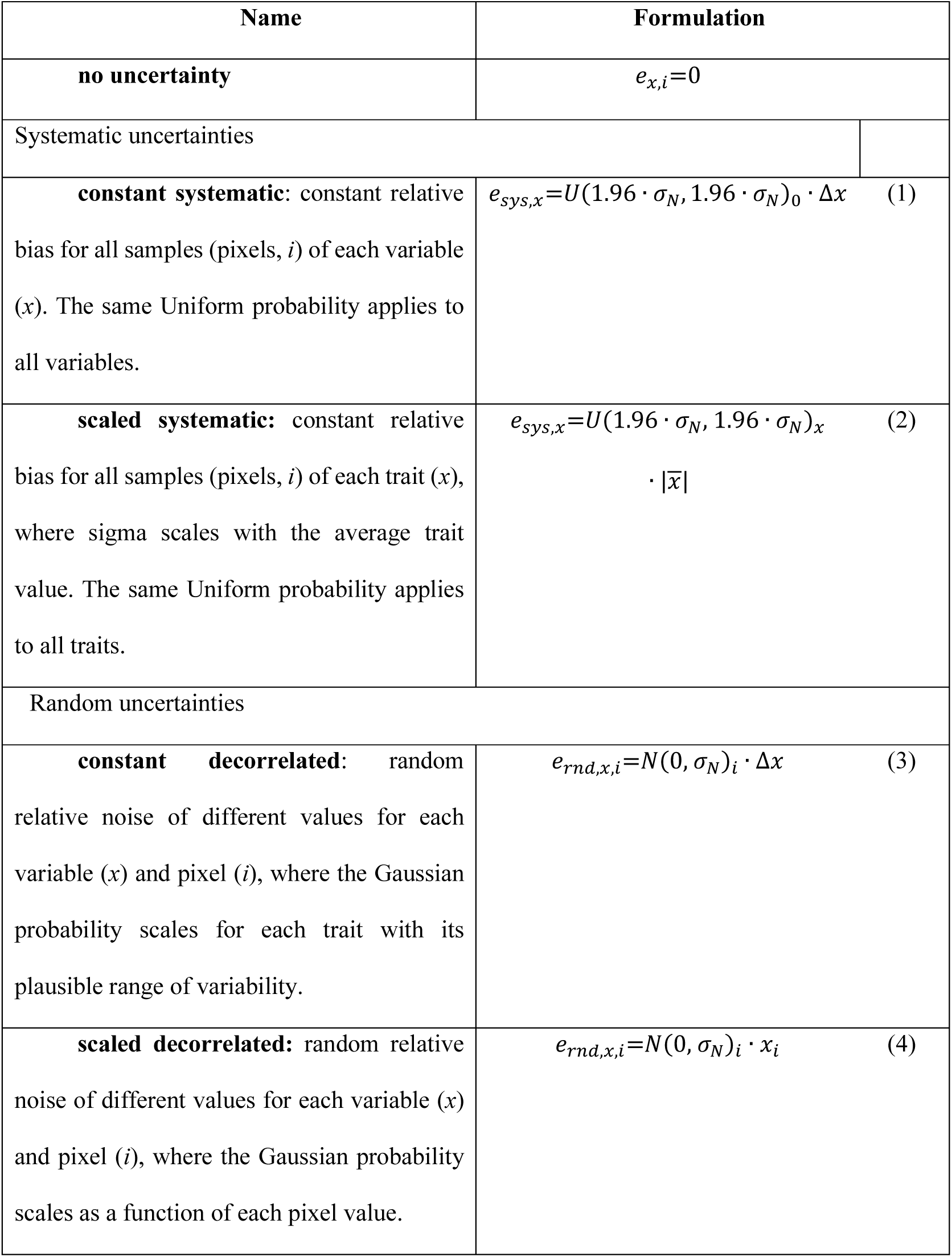
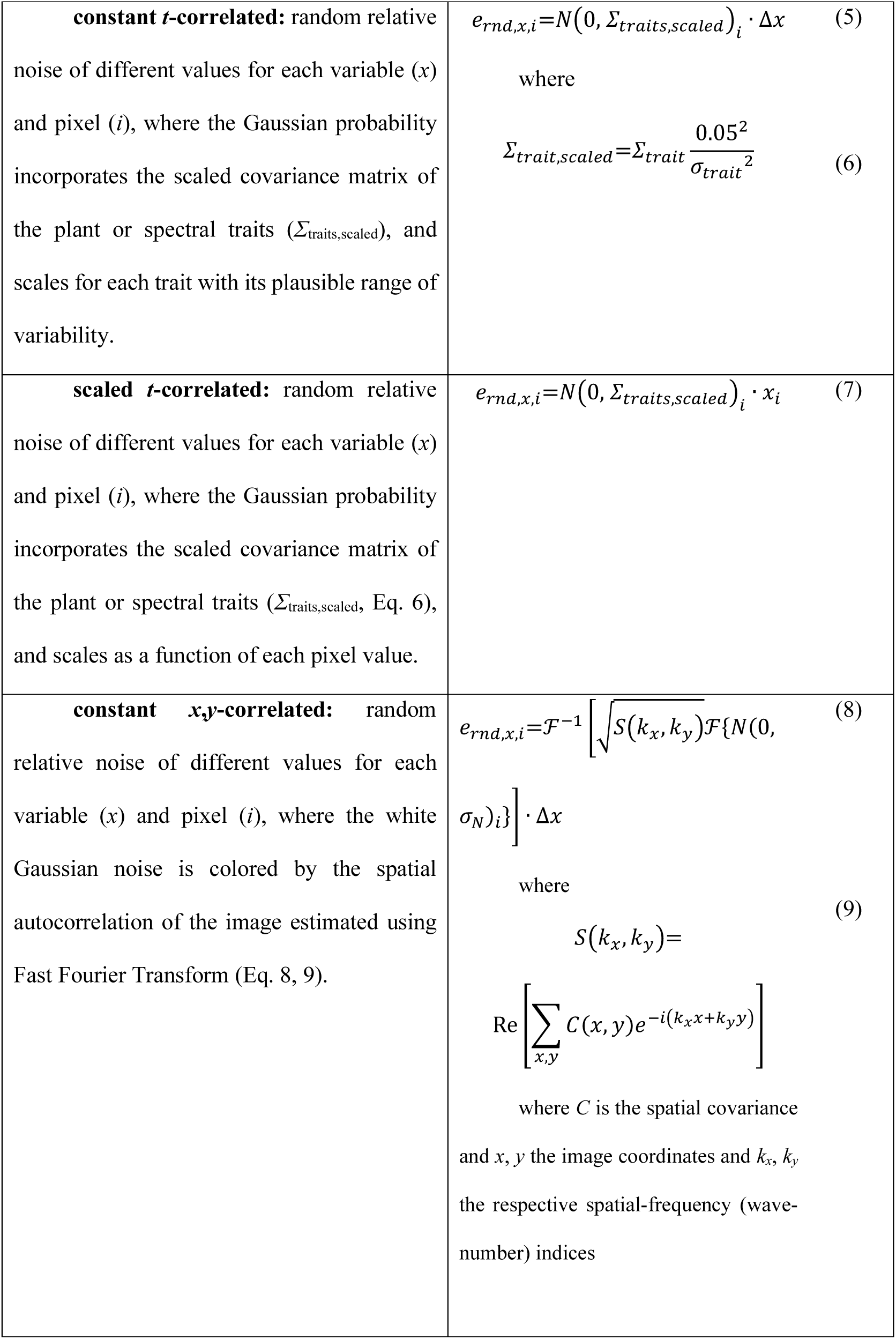
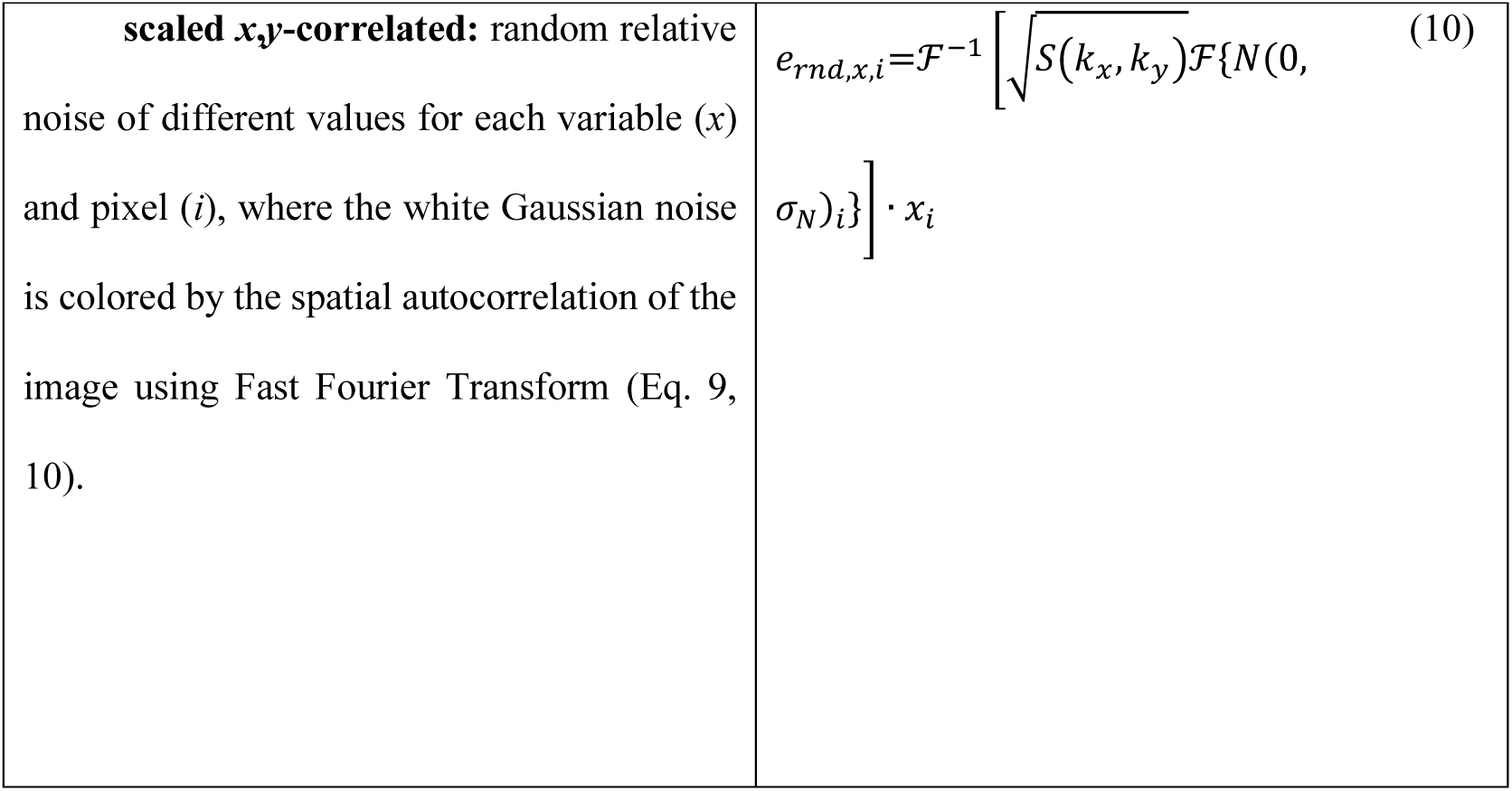
Definition and formulation of uncertainty types.

Errors were first defined in relative terms. Thus, we set the standard deviation of the Gaussian distribution to 5 % (*N*(0, *σ*_*N*_), where *σ*_*N*_ = 0.05), a value in accordance with the mission requirements for the surface reflectance product of DESIS (de los Reyes et al., 2020) and with results from other spaceborne imaging spectrometers, such as EnMAP and PRISMA, which exhibit absolute reflectance differences of approximately 2–7% relative to ground validation data (Brell et al., 2026; Cogliati et al., 2021). Then, we scaled *σ*_*N*_ (representing the 68 % of the confidence range in the Gaussian distribution) as 1.96 × *σ*_*N*_ to define a Uniform distribution of the same coverage (i.e., *U*(-1.96 × *σ*_*N*_, 1.96 × *σ*_*N*_)) (Eq. 1-2).

Furthermore, “random” uncertainties were simulated both fully “decorrelated” (Eq. 3-4) and correlated in two different ways (Eq. 5-10). First, we imposed trait correlation (“*t*-correlated”), using the covariance of the spectral or plant traits of each scene as a proxy for the uncertainty covariance. We scaled each covariance matrix by computing the correlation matrix and multiplying it by the squared 5% relative uncertainty (Eq. 6). Second, we imposed spatial autocorrelation (“*x*,*y*-correlated”) (Wood and Chan, 1994). To do so, we standardized each trait or spectral band and averaged the trait 3D cube into a 2D image. Then, we computed the covariance from single spatial lags (Eq. 11) and applied circulant embedding (Dietrich and Newsam, 1997) to avoid the artificial periodic boundary condition introduced by the ordinary Fast Fourier Transform. Then, we used the Wiener-Khintchine theorem (Khintchine, 1934; Norbert Wiener, 1930), which states that the power spectral density of a wide-sense-stationary random process equals the Fourier Transform of that process’s autocorrelation function. Therefore, we extracted the power spectral density eigenvalues from the embedded covariance matrix (Eq. 9) and used them to “color” the (decorrelated) white noise initially generated (Eq. 8, 10). Finally, we preserved the scale of the original noise by normalizing it with the ratio of the original to the correlated standard deviations.

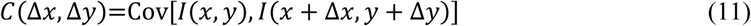

where *I* is the averaged 2D image of plant traits or reflectance factors, Δ*x* and Δ*y* are the discrete lags (in pixels) used to compute the autocovariance function (*C*).

For each realization, a single value from the uniform distribution was drawn to represent the systematic error across all samples (i.e., pixels); whereas for random uncertainties, different error values for each sample were drawn from the Gaussian distribution. We also considered two additional types of uncertainty, those independent of the PT or *R* magnitude (“constant”) and those that scale with them (“scaled”). In the first case, we multiplied the relative error value drawn from the PDF by the range of values of each trait (PT or *R*) in each scene (Δ*x*). In the second case, we multiplied the relative error value drawn from the PDF by the PT or *R* magnitude (or the mean for the “systematic” uncertainty). Crossing constant/scaled and systematic/random (decorrelated, *t*-correlated, and *x*,*y*-correlated) uncertainties (Fig. 1b), we simulated eight different uncertainty types (Table 3).

Finally, to ensure the comparability between the absolute scaled and constant uncertainties, constant uncertainties were rescaled by using the standard deviation of the errors of the analogous scaled uncertainty (Eq. 12), so that the root mean squared error (RMSE) of each variable was equivalent for each scene:

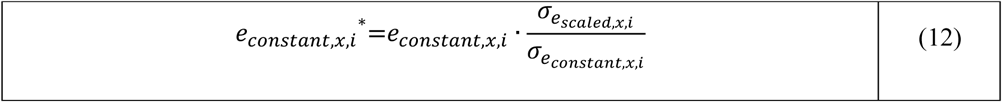

### 2.3 Functional diversity estimation

For each plant or spectral trait (Table 2), we computed the functional diversity metric Rao’s quadratic entropy (*Q*_Rao_, Eq. 13) using the pyGNDiv package (Pacheco-Labrador et al., 2023). pyGNDiv normalizes the dissimilarity metric with respect to trait dimensionality, making the metrics across dataset types directly comparable (i.e., at the 1:1 line). By default, pyGNDiv applies standardization, dimensionality reduction with principal component analysis, and dissimilarity normalization when using PT, *R*, SVI_3_, and OT. In this case, the dissimilarity (Euclidean distance) is computed using the principal components selected to retain 98% of the variance. However, NDVI and NIR_v_ proxies feature a single variable and underwent only standardization. In all cases, we computed *Q*_Rao_ using a small (3-by-3) moving window, provided *Q*_Rao_ tends to saturate and converge as the window size increases (Pacheco-Labrador et al., 2026). This resulted in 1,600 *Q*_Rao_ values per PT map or remote sensing proxy.

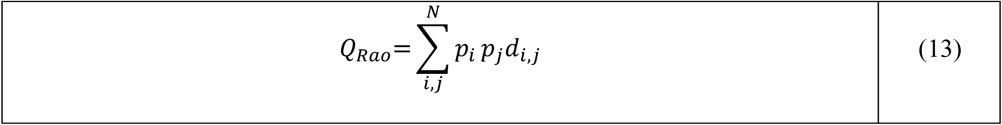

where *p* is the relative abundance of the samples *i* and *j*, and *d*_ij_ is the dissimilarity between the traits of both samples, in this case, the Euclidean distance normalized for dimensionality (Eq. 14).

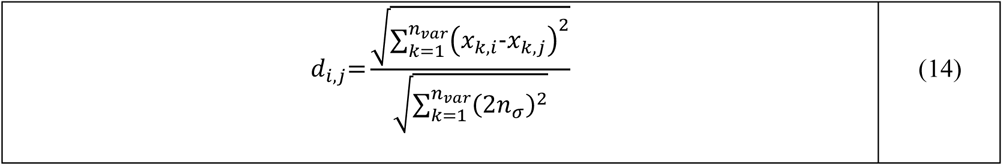

where *n*_var_ is the number of traits of the dataset, *x* is the value of the principal component *k* or a unidimensional standardized spectral index for the samples *i* and *j*, and *n_σ_* is a distance defined in standard deviations in the z-score scale (by default, 6).

### 2.4 Uncertainty impact evaluation

We assessed the effect of each uncertainty type by regressing the *Q*_Rao_ computed from the synthetic plant traits maps (*Q*_Rao,PT_) against that from the remote sensing proxies (*Q*_Rao,RS_), and computed a set of metrics of performance, including the coefficient of determination (*R*^2^), Pearson’s correlation coefficient (*r*_Pearson_), relative root mean squared error (RRMSE, [%]), mean absolute error (MAE, [-]), and the slope (*b*_1_) and bias (*b*_0_) of the predicted (*Q*_Rao,RS_) - observed (*Q*_Rao,PT_) relationship (Fig. 1d). In both cases we considered the “true error”, this is, we compared *Q*_Rao,RS_ under uncertainty vs. *Q*_Rao,PT_ under no uncertainty. However, we additionally considered the “apparent error”, where *Q*_Rao,PT_ also incorporates uncertainty (Widlowski, 2015), to identify enhancing or diminishing effects.

The evaluation took place at two scales: “between” and “within” sites. In the first case, we compared the mean *Q*_Rao,PT_ and *Q*_Rao,RS_ for each scene and date (720 pairs of data in total) simultaneously to assess the impact of uncertainty on PFD characterization at regional or global scales (“between” sites). For the “within” scenes case, we repeated the evaluation for each scene and date independently, comparing the 1,600 pairs of *Q*_Rao,PT_ and *Q*_Rao,RS_ values of each scene, and examined the resulting distributions of the abovementioned statistics (180 scenes × 4 dates). This second analysis assesses how uncertainty affects PFD characterization within small study areas.

## 3. RESULTS

### 3.1 Uncertainty simulation

Fig. 3 exemplifies how the absolute (*e*_abs_, Fig. 3a) and relative (*e*_rel_, Fig. 3b) errors are generated over a synthetic data cube of uniformly distributed values ranging from 0 to 1, sliced into 19 layers at 0.05-step intervals and 0.1 width (Fig. S1). Fig. 3a also shows the standard deviation of the simulated errors (*σ_e_*) and the RMSE relative to the values simulated without uncertainty. As shown, the RMSEs for constant and scaled uncertainties are the same. Also, constant uncertainties are independent of the trait magnitude (*x*-axis) in absolute terms (Fig. 3a); whereas relative uncertainty is larger for low values (Fig. 3b). Scaled systematic also produces larger relative uncertainties at low trait values, whereas scaled random (decorrelated and correlated) relative uncertainties do not, except for *x*,*y*-correlated. In this case, as the traits were generated independently, decorrelated and *t*-correlated uncertainties behave similarly. However, this may not hold for autocorrelated variables. The spectral reflectance factors feature high spectral correlation, making correlated errors more similar to the systematic ones (Fig. S2). The covariance between plant traits is nonetheless lower, becoming the character of the error trait-specific (i.e., *C*_ab_, Fig. S3). Spatial correlation generated spatial patterns that diminished from clustered to even spectral patterns (Fig. S4).

**Figure 3.**
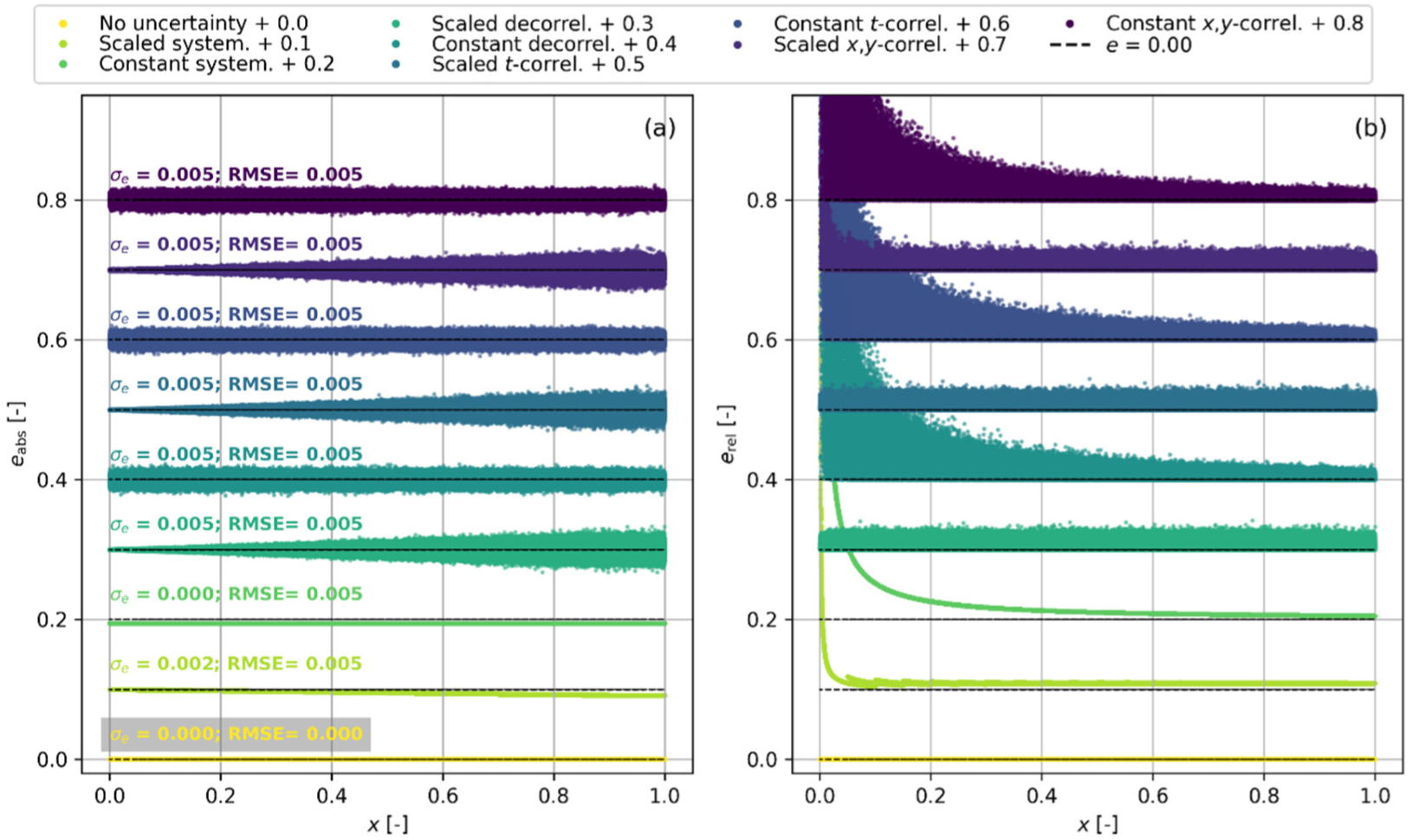
Absolute (a) and relative (b) errors (*e*) simulated for different uncertainty types over a synthetic dataset ranging between 0 and 1. For each type, “constant” errors are scaled to meet the same “scaled” root mean squared error with respect to the original value (RMSE). The standard deviation of the errors applied (*σ_e_*) is also shown.

### 3.2 Uncertainty propagation to diversity estimation: Between-site comparison

Fig. 4 presents the predicted-observed relationships between mean scene *Q*_Rao_ computed from plant traits (*Q*_Rao,PT_) without uncertainty and each remote sensing proxy under each uncertainty type (“true error”, “between sites”; 720 data points). PT diversity (*Q*_Rao,PT_) was insensitive to systematic uncertainties (Fig. 4a); the small discrepancy (RRMSE ≤ 0.14%) likely resulted from changes in the number of principal components selected in a few cases. Thus, systematic uncertainties in PT did not alter the relationship with remote sensing estimates of *Q*_Rao_ (Fig. S5b-f). However, random (decorrelated and correlated) uncertainties led to slight overestimation, with a larger impact at the lowest diversity values.

**Figure 4.**
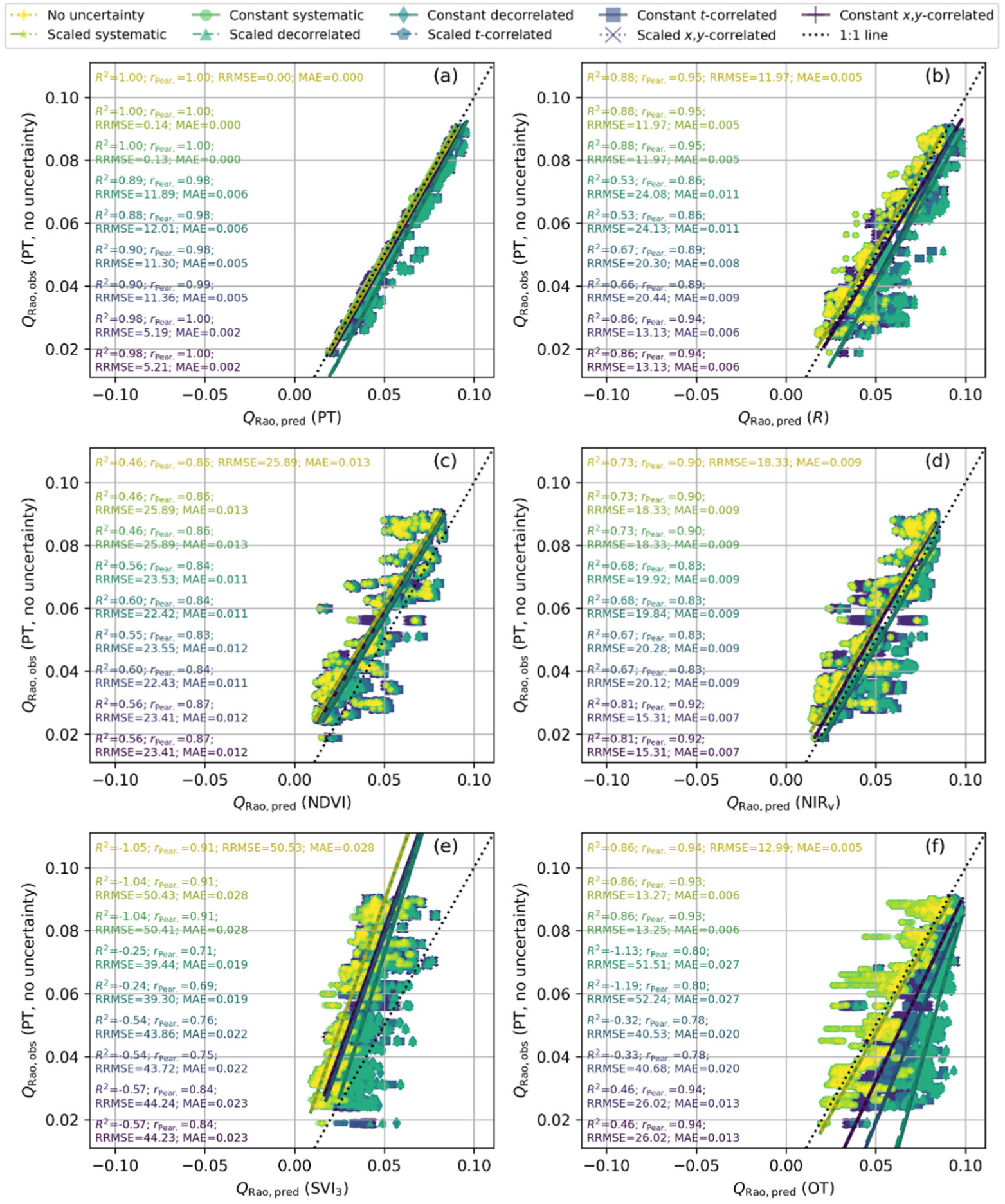
Comparison of Rao’s quadratic entropy (*Q*_Rao_) computed from plant traits without uncertainty and from plant traits (a), reflectance factors (b), normalized difference vegetation index (c), near-infrared reflectance of vegetation (d), a cube of three spectral indices (e), and optical traits (f) featuring different data uncertainty types (“true error”).

*Q*_Rao_ computed from reflectance factors (*Q*_Rao,*R*_) without uncertainty closely matched *Q*_Rao,PT_ (Fig. 4b) and was insensitive to systematic uncertainty. Contrarily, random (decorrelated and correlated) uncertainties led to overestimation, with each pair of constant and scaled uncertainties behaving almost identically. Trait and spatial correlation uncertainties diminished the overestimation of high and low *Q*_Rao,PT_ values, respectively.

The unidimensional spectral vegetation indices (NDVI (Fig. 4c) and NIR_v_ (Fig. 4d)) underestimated *Q*_Rao,PT_, and were barely affected by systematic uncertainties; whereas random uncertainties increased *Q*_Rao_ values, partly compensating for the original underestimation (i.e., reducing RRMSE). For NDVI and NIR_v_, random uncertainties behaved very similarly and approximated the predicted-observed relationship slightly above the 1:1 line (Fig. 4c,d).

SVI_3_, being a more complex byproduct of *R*, underestimated *Q*_Rao,PT_, particularly the large values, and under all uncertainty types (Fig. 4e). There were minimal differences between constant and scaled uncertainty types, but errors showed larger variability for the different bootstrap realizations (along the *x*-axis) than the variables already analyzed (Fig. 4a-d). The systematic uncertainties produced the smallest effects with respect to the original relationship, sustaining the largest biases and the largest *r*_Pearson_. Contrarily, random uncertainties led to a relatively constant overestimation, minimizing errors but also *r*_Pearson_ correlation. The correlated uncertainties produced smaller changes for the lowest *Q*_Rao,PT_ values and intermediate performance.

Finally, OT produced values close to *Q*_Rao,PT_ (Fig. 4f), with performance comparable to *R* (Fig. 4b) under no or systematic uncertainty. However, random and, in particular, decorrelated uncertainties led to the largest overestimation across all remote sensing proxies, peaking at low *Q*_Rao,PT_ values, with a similar effect for constant and scaled types. Trait and then spatial correlation in random uncertainty reduced its impact. OT also showed the greatest variability across bootstrap realizations (along the *x*-axis).

Furthermore, the “apparent error” assessment showed that PT random uncertainties (correlated or not) can diminish the effect of remote-sensing overestimation (Fig. S5), reducing biases and changes in the slope of the predicted-observed relationship. The improvement in the statistics was most notable for *R* (Fig. 4b vs. Fig. S5b) and OT (Fig. 4f vs. Fig. S5f), followed by NIR_v_ (Fig. 4d vs. Fig. S5d), where RRMSE increased sometimes. Statistics worsened for NDVI and SVI_3_ (Fig. 4c,e vs. Fig. S5c,e). However, these statistical improvements are spurious, as both *Q*_Rao_ estimates are indeed biased with respect to the true values.

### 3.3 Uncertainty propagation to diversity estimation: Within-site comparison

Fig. 5 presents statistics assessing the impact of different types of data uncertainty on PFD estimation (*Q*_Rao,PT_) using various remote sensing proxies (*Q*_Rao,RS_; *x*-axis) “within sites”. Each of the 720 data points corresponds to a statistic comparing the 1,600 *Q*_Rao_ computed in each trait map or remote sensing image (“true error”). Analogously, Fig. 6 shows the corresponding difference (*Δ*) between the statistics obtained under each uncertainty type and the reference case under no uncertainty. Overall, *Q*_Rao_ computed from OT and SVI_3_ were the most affected, and constant decorrelated uncertainty produced the strongest impacts, followed by *t*-correlated.

**Figure 5.**
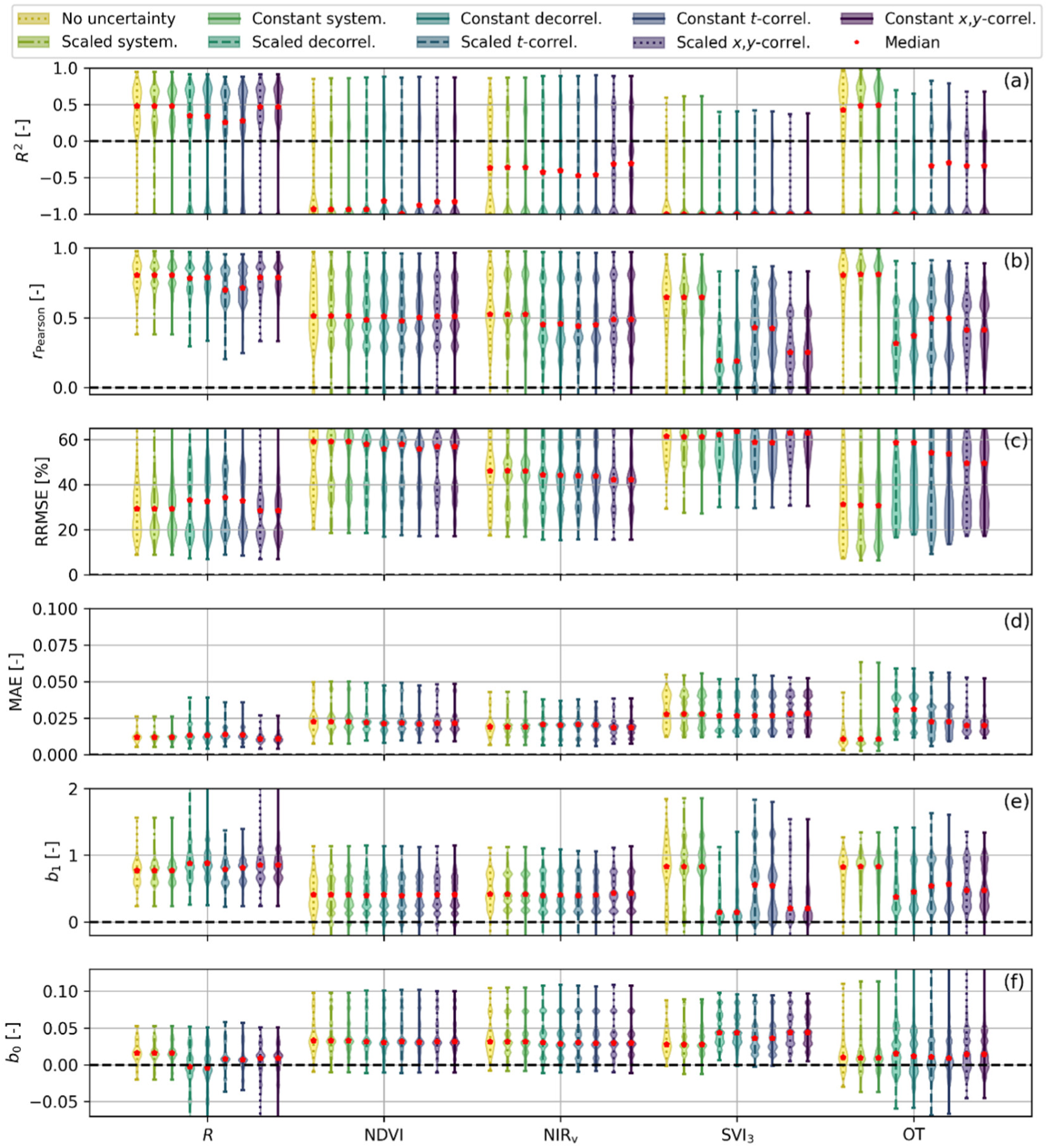
“True error” statistics of the performance in the estimation of plant functional diversity for the different types of uncertainty: (a) coefficient of determination, (b) Pearson correlation coefficient, (c) relative root mean squared error, (d) mean absolute error, slope (e) and offset (f) of the predicted-observed relationship.

**Figure 6.**
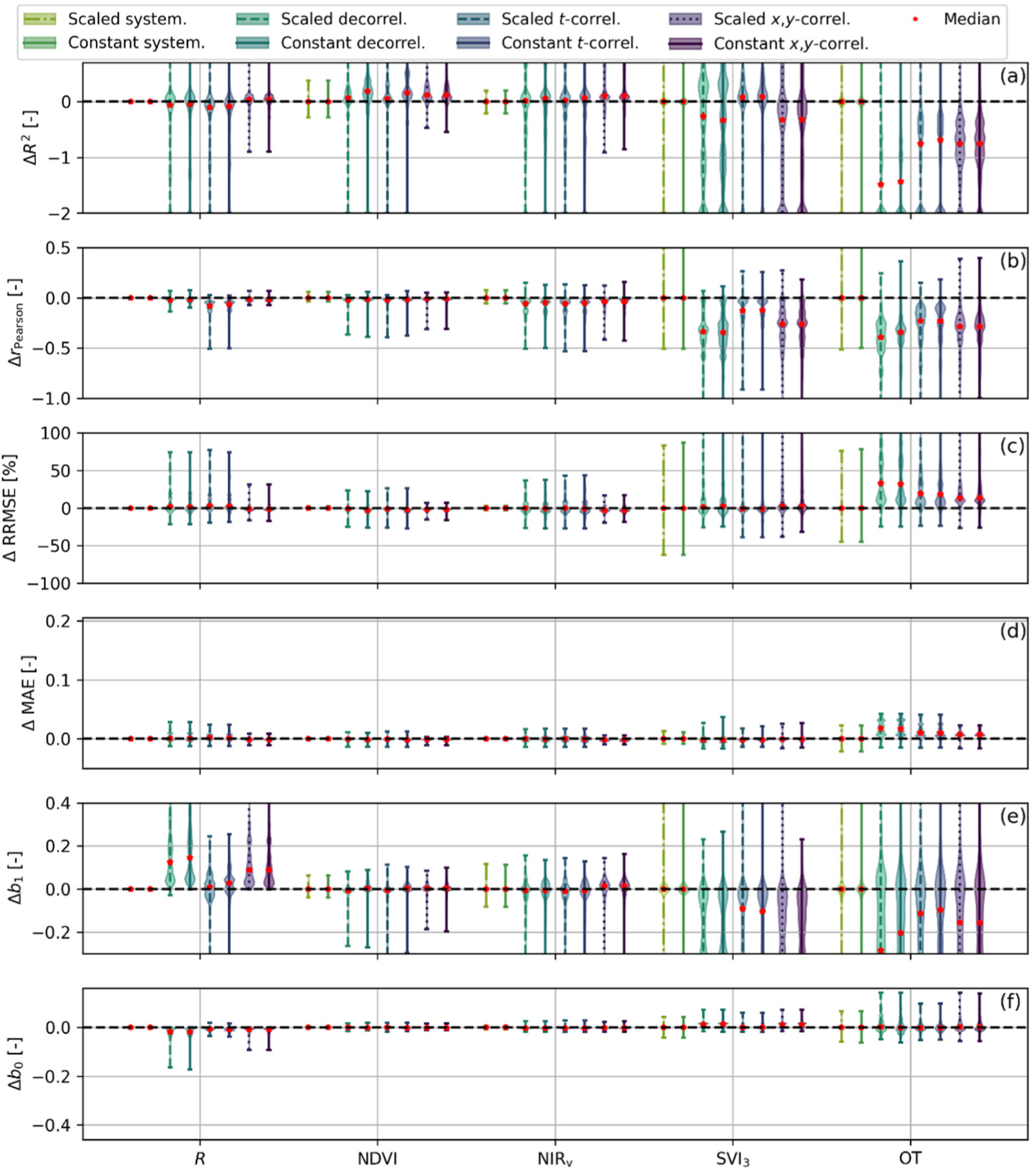
Difference of “true error” statistics of the performance in the estimation of plant functional diversity with the case “no uncertainty” for the different types of uncertainty: (a) coefficient of determination, (b) Pearson correlation coefficient, (c) relative root mean squared error, (d) mean absolute error, slope (e), and offset (f) of the predicted-observed relationship.

Under no uncertainty, *R* and OT achieved the best estimates of PFD (i.e., the largest correlations, the lowest errors, and predicted-observed relationships closest to the 1:1 line) (Fig. 5). Among the remaining remote sensing proxies, NIR_v_ achieved the most accurate estimates (the largest *R*^2^ (Fig. 5a) and the lowest RRMSE and MAE (Fig. 5c-d)); SVI_3_ showed the strongest *r*_Pearson_ (Fig. 5b), but also the largest RRMSE and MAE (Fig. 5c-d)); whereas NDVI behaved similarly to NIR_v_, but with larger errors and lower *R*^2^.

Overall, constant and scaled uncertainties had similar effects (Fig. 5,6), whereas systematic uncertainties had almost negligible effects. The median difference between statistics deviated from 0 (meaning no change in average) only for SVI_3_ and OT (and *R* for *b*_1_), where decorrelated (random) uncertainties had the largest impact on performance (Fig. 6a-d). The impact of decorrelated and *x*,*y*-correlated uncertainties was similar for NDVI and NIR_v_ (Fig. 6a-d), which were most affected by *t*-correlated uncertainties.

Most often, random uncertainties increased the offset (*b*_0_, Fig. 6f) and reduced the slope (*b*_1_, Fig. 6e) of the predicted-observed relationship, inducing larger changes than systematic uncertainties. For *R*, the impact went in the opposite direction, approximating the median relationship to the 1:1 line. All uncertainties had minimal effects on the NDVI and NIR_v_ predicted-observed relationship (Fig. 6e-f).

Finally, when considering the “apparent errors”, we found that the uncertainty in PT modified the impact of random uncertainty types on performance (Fig. S6-S7), slightly reducing correlations (Fig. S6a-b and S7a-b), but also RMSE and slope (*b*_1_) (Fig. S6c,e and S7c,e), while increasing the offset (Fig. S6f and S7f), and increasing MAE for NDVI and SVI_3_ (Fig. S6d and S7d).

## 4. DISCUSSION

The effect of uncertainty in the estimation of PFD (i.e., *Q*_Rao_) from remote sensing depends on the type of data uncertainty, the remote sensing proxy used to compute *Q*_Rao_, and the preprocessing of both the remote sensing proxies and the in-situ measured plant functional traits. Overall, systematic errors (i.e., biases) had a smaller impact on PFD estimation than random decorrelated and correlated errors (i.e., noise, Fig. 4-6), partly because of the preprocessing used to compute the functional diversity metric (i.e., *Q*_Rao_). For random uncertainties, trait correlation led to a stronger impact on PFD estimates than spatial correlation. The impact of uncertainties also increased with the complexity of the remote sensing proxy used to estimate PFD.

### 4.1 Uncertainty correction via preprocessing

Our results suggest that standardizing in-situ plant trait measurements and reflectance factors before computing the diversity metric cancels out or reduces the impact of systematic errors. Mean subtraction removes trait-specific biases before computing the PFD metric (in this case, *Q*_Rao_). Standardization is necessary for two reasons: a) removing the differences in magnitude between traits or remote sensing proxies so that their variabilities are not spuriously weighted (Magneville et al., 2022), and b) the application of dimensionality reduction, given that collinearity can spuriously modify diversity (Laliberté and Legendre, 2010). However, these steps can be avoided in ecology in favor of selecting traits considered relevant or prioritizing interpretability (Zhu et al., 2017), and they are not always applied in remote sensing. For example, Helfenstein et al. (2022) and Schneider et al. (2017) applied normalization instead. Additional attention could be paid to the distribution of the traits and abundances, as it is known that this can affect the estimation of PFD, particularly when data are missing (Májeková et al., 2016); although Normality is not a requirement for the computation of diversity metrics (de Bello et al., 2010; Laliberté and Legendre, 2010; Ricotta and Szeidl, 2006) and standardization might be more relevant (Magneville et al., 2022). Using simulations, Pacheco-Labrador et al. (2022) reported that omitting these steps reduced the correlation between functional diversity metrics derived from plant traits and remote sensing proxies. We repeated our analyses without applying standardization or dimensionality reduction to the traits prior to the computation of *Q*_Rao_ and observed an overall decrease in performance, particularly “between sites” and for variables with the largest dimensionality (compare Fig. 4-6 vs. Fig. S8-S10), i.e., the hyperspectral reflectance factors (197 spectral traits, i.e., bands) and the optical traits (12 estimated plant traits).

Our results suggest that this mathematical abstraction (standardization and dimensionality reduction) may not only be a necessary step to link field-based and remote sensing-based estimates of PFD, but also a step that reduces their uncertainty. The application of this result to remote sensing is straightforward: functional diversity estimation from remote sensing would not require image-correction processes applied uniformly across each spectral band (e.g., empirical line correction, often applied to drone or airborne imagery to transform at-sensor radiance or raw digital numbers into *R*) (Smith and Milton, 1999).

Additionally, dimensionality reduction might help to attenuate random errors, as the fraction of variance retained could be adjusted based on the magnitude of the noise, if it were known (e.g., via uncertainty propagation). We did not assess the potential of this correction, but algorithms such as PCA, used in this study, are often recommended for denoising multidimensional variables (Corner et al., 2000).

### 4.2 The impact of uncertainty on plant traits and remote sensing proxies

Remote sensing characterizes vegetation using bottom-of-atmosphere spectral signals that exclude atmospheric contributions. Like most studies, ours departed from reflectance factors and tested variables derived from them at different levels of complexity (spectral indices and optical traits). Our results suggest that the impact of uncertainty depends on the proxies used to estimate PFD, and more importantly, that it increases with the degree of elaboration from the primary variable, in this case, the reflectance factors. While our results are limited to the specific remote sensing proxies we used, the pattern we found is explainable and may be generalized. The reflectance factor is the primary source of information, as the fraction of radiation reflected by the surface is determined by its structural and biochemical properties and relates nonlinearly to these variables (including plant functional traits) (Jacquemoud and Baret, 1990; Verhoef, 1984). While standardization can remove systematic uncertainties from *R*, spectral index formulation might partially mitigate this correction by propagating such biases inhomogeneously across samples or pixels, and some might even inflate the impact of certain uncertainty types. Uncertainty propagation to optical traits strongly depends on the retrieval method used, but estimating traits with overlapping absorption features or weak effects on reflectance factors can be inherently uncertain (e.g., Féret et al. (2019)). Our results show that *R*, as an estimator of *Q*_Rao,PT_ is more robust to uncertainty than simple SVIs (i.e., NIR_v_ and NDVI), and these are more robust than more complex combinations of indices (e.g., SVI_3_) or OT.

However, *R* is not always a preferred choice for estimating PFD. For example, remote sensing is expected to contribute to global biodiversity mapping through the Essential Biodiversity Variables, which are primarily SVIs, OT, or other reflectance-based estimates (Skidmore et al., 2021). Furthermore, *R*, particularly hyperspectral, requires more computational resources to compute PFD metrics than SVIs or lower-dimensional OT proxies. In addition, diversity computed from spectral variables (*R* and SVIs) is highly sensitive to background effects (Gholizadeh et al., 2018; Wang et al., 2018b), a sensitivity OT can alleviate (Pacheco-Labrador et al., 2026). On the other hand, OT estimation also introduces uncertainties that affect PFD estimation. Beyond the methods and variables here tested, additional preprocessing approaches have been proposed to address various sources of uncertainty affecting PFD estimation (e.g., shadow removal) (Perrone et al., 2024; Rossi and Gholizadeh, 2023; Wallis et al., 2025); for which the impact of uncertainty propagation could also be analyzed.

In this context of trade-offs and possible choices, dimensionality-normalized diversity metrics, which offer one-to-one comparability across different PFD estimates (e.g., Pacheco-Labrador et al. (2023)), would facilitate sanity checks and quality controls by comparison of estimates from different remote sensing proxies (e.g., *R* and OT). Such normalization not only allows for comparison between remote sensing-based functional diversity estimates but also, when available, for comparison with field measures of plant traits for validation. This exercise may be relevant, particularly when no field data have been collected using dedicated protocols to validate remote sensing estimates of functional diversity (e.g., Hauser et al. (2021)).

We also found that random uncertainties in the plant traits counteracted the performance loss induced by the same uncertainty type in the remote sensing proxies, or even improved performance relative to the uncertainty-free case. However, statistical improvements are proxy-and extent-dependent (“between” and “within” sites) (Fig. 4-6 vs. S5-S7); their interpretation is not straightforward and may reflect spurious effects. Under uncertainty, both diversity estimates (from in-situ data and remote sensing proxies) are biased (Fig. 4a), yet in-situ data is considered the reference estimate of PFD, independently of its uncertainty. Thus, quantifying its uncertainty (rarely done) may be relevant for assessing the value of ground measurements in validating remote sensing estimates of PFD.

Spatial and, in particular, trait error correlation reduced the impact of random uncertainties simulated as white Gaussian noise. Error correlation can increase, reduce, or cancel the effect of different errors through error propagation, depending on the sign of the correlation and the model sensitivities (JCGM, 2008; Zhu et al., 2018). Our simulations showed attenuation of uncertainties, matching previous work where spectral or spatial autocorrelation of model inputs reduced uncertainty (Zhang et al., 2024); which suggests, accounting for autocorrelation is necessary to prevent random error uncertainty overestimation (Rossi et al., in press). However, we cannot assure that such an effect applies generally; for example, BOSSE scenes’ autocorrelation is null or positive, but not significantly negative (Pacheco-Labrador et al., 2026), which matches common findings in observations (Karasiak et al., 2019; Li and Leighton, 1992), but leaves such a case untested. Consequently, while uncertainty may most often attenuate the impact of correlation, we must stress the need for careful uncertainty propagation.

Beyond uncertainties, the different remote sensing proxies showed varying capabilities for estimating PFD, which was computed from a set of foliar and structural traits, with varying impacts on the reflectance factors. In this regard, the systematic underestimation of NDVI and NIR_v_, and the lower performance of SVI_3_ can be explained by this comparison, as they do not capture all the diversity explained by the simulated PT. Indices like SVI_3_ are supposed to be mostly sensitive to foliar traits (Schneider et al., 2017), whereas structural variables also explain an important fraction of the spectral variability (Hauser et al., 2021b; Rossi et al., 2021). In this regard, simulations such as those enabled by BOSSE can support the optimization or analysis of relationships between functional diversity metrics derived from remote sensing and specific sets of plant traits.

### 4.3 Beyond simulations and potential limitations

BOSSE provided a suitable framework for assessing the effects of different types of uncertainty on estimating PFD (i.e., *Q*_Rao_) from remote sensing datasets. However, limitations in the simulated scenes and uncertainties must be considered, so the conclusions are guidelines rather than strict rules to follow blindly. BOSSE represents simple canopies (vertically homogeneous) and therefore does not allow assessment of uncertainties introduced by trait vertical heterogeneity or the presence of understory plants. Also, despite simulating six statistical types of uncertainty, our simulations did not include relevant sources of uncertainty such as spectral mixture or the presence of shadows produced by crown light occlusion, among others, some of which are already known to have strong effects on the estimation of PFD (Pacheco-Labrador et al., 2026; Wallis et al., 2025; Wang et al., 2018a). Still, these could operate as both systematic and random uncertainties, as simulated in this study, whereas approaches to process or filter the data can attenuate or inflate the effect of these sources of uncertainty. Also, the uncertainty incorporated by the retrieval of optical traits strongly depends on the methods and radiative transfer models used; our retrieved OTs presented relatively large errors, even in the absence of uncertainties in *R* (Fig. S11), which increased the *Q*_Rao_ estimation error with respect to other remote sensing proxies (e.g., Fig. 4,5). Beyond simulations, it is still necessary to produce observational datasets sufficiently complete to further explore, optimize, or validate methods for estimating PFD from remote sensing. These should include sets of measured plant traits (or biophysical variables) deemed sufficient to explain canopy spectral signals (both foliar and structural). Here, sensitivity analyses of radiative transfer models could provide insight into the most relevant plant traits and the amount of reflectance variance a given set could potentially explain (e.g., Verrelst et al. (2015)). Furthermore, these should be sampled using *ad hoc* protocols that allow validation of remote sensing estimates of PFD (e.g., Hauser et al. (2021a)) and be sufficiently intensive to minimize representation uncertainty (Albert et al., 2010). At the same time, remote sensing data should feature high spatial and spectral resolutions (e.g., Cimoli et al.(2024)) to provide complete information on plant properties and ensure the matching between the image and the plant individuals, minimizing thus the impact of spatial mismatch and spectral mixture (Ludwig et al., 2024; Pacheco-Labrador et al., 2026; Wang et al., 2018b, 2018a), which might also be challenging. Finally, both plant traits and remote sensing imagery should include uncertainty budgets. These would need to account for measurement, but also representation and model uncertainty (e.g., OT retrieval), and separate the systematic and random components and their correlation, since, as we have shown, biases can have null or minimal impact for certain remote sensing proxies, and correlation can diminish error propagation to PFD. Such a dataset (or rather a collection of several, encompassing a representative range of diversity ranges and vegetation types) is highly demanding, which might make it unlikely. Intermediate steps, such as estimation of optical trait model uncertainties (e.g., Wang et al. (2019)) or datasets partially meeting these requirements (e.g., Hauser et al. (2021b)), might provide helpful, incomplete, but complementary insights, or enable the assessment of very specific methodological questions (e.g., Hauser et al. (2021b)). At the same time, simulations can also be made more complex, e.g., incorporating models describing geometrical components or vertical trait heterogeneity (Gastellu-Etchegorry et al., 2004; Henniger et al., 2023). In an operational context, it is critical that remote sensing products incorporate uncertainty estimates for each variable (e.g., uncertainty cubes) (Rossi et al., in press), so that the reliability of different variables or areas of the image can be evaluated and these disregarded if necessary.

## 5. CONCLUSIONS

The Biodiversity Observing System Simulation Experiment helped assess the impact of different types of uncertainty in estimating PFD from remote sensing. We found that preprocessing plant and spectral trait datasets removed or reduced the impact of systematic biases; however, this correction loses effectiveness as the remote sensing variables derived from reflectance become increasingly complex. Error correlation generally reduced the impact of random errors, which were also attenuated by the presence of the same type of error in the plant trait measurements. We also found that uncertainties can spuriously correct for inherent biases between diversity values estimated from in-situ plant trait measurements and different remote sensing proxies. Despite the limitations of simulations, our results suggest that uncertainty propagation in plant functional diversity estimation must be accounted for differently from that in the more traditional problem of estimating the “mean” trait value. This understanding is relevant for the design and optimization of new methodologies and protocols.

In the future, more extensive datasets and complex modeling exercises should provide more robust and comprehensive insights into how different sources of uncertainty affect the estimation of plant functional diversity from space, helping define the limitations of this technology and quantify the uncertainty of dedicated remote sensing products.

## ACKNOWLEDGEMENTS

JPL acknowledges the project “Integrated Observing Systems and Simulation Experiments to Analyze Biodiversity-Ecosystem Function Relationships in Savanna Ecosystems” PID2023-151046NB-I00 funded by MCIU/ AEI / 10.13039/501100011033 / FEDER, UE). CR and MJS acknowledge the Horizon Europe OBSGESSION project (Grant Agreement No. 101134954)

## Data availability

The Python code used to run the simulations and figures of this manuscript can be found in Zenodo: https://doi.org/10.5281/zenodo.22872340 (Pacheco-Labrador, 2026). The Biodiversity Observing System Simulation Experiment (BOSSE v1.0) code is available on GitHub: https://github.com/JavierPachecoLabrador/pyBOSSE, and the additional ERA5Land Meteorological time series are available in Zenodo: https://doi.org/10.5281/zenodo.14717038 (Pacheco-Labrador et al., 2025). The pyGNDiv package used to compute normalized functional diversity metrics can be found on GitHub: https://github.com/JavierPachecoLabrador/pyGNDiv-master.

## REFERENCES

Albert, C.H., Yoccoz, N.G., Edwards Jr, T.C., Graham, C.H., Zimmermann, N.E., Thuiller, W., 2010. Sampling in ecology and evolution – bridging the gap between theory and practice. Ecography 33, 1028–1037. 10.1111/j.1600-0587.2010.06421.x

Asner, G.P., Martin, R.E., Knapp, D.E., Tupayachi, R., Anderson, C.B., Sinca, F., Vaughn, N.R., Llactayo, W., 2017. Airborne laser-guided imaging spectroscopy to map forest trait diversity and guide conservation. Science 355, 385. 10.1126/science.aaj1987

Bachmann, M., Makarau, A., Segl, K., Richter, R., 2015. Estimating the Influence of Spectral and Radiometric Calibration Uncertainties on EnMAP Data Products—Examples for Ground Reflectance Retrieval and Vegetation Indices. Remote Sensing 7, 10689–10714.

Badgley, G., Field, C.B., Berry, J.A., 2017. Canopy near-infrared reflectance and terrestrial photosynthesis. Science Advances 3, e1602244. 10.1126/sciadv.1602244

Beccari, E., Pérez Carmona, C., Tordoni, E., Petruzzellis, F., Martinucci, D., Casagrande, G., Pavanetto, N., Rocchini, D., D’Antraccoli, M., Ciccarelli, D., Bacaro, G., 2024. Plant spectral diversity from high-resolution multispectral imagery detects functional diversity patterns in coastal dune communities. Journal of Vegetation Science 35, e13239. 10.1111/jvs.13239

Botta-Dukát, Z., 2005. Rao’s quadratic entropy as a measure of functional diversity based on multiple traits. Journal of Vegetation Science 16, 533–540. 10.1111/j.1654-1103.2005.tb02393.x

Brell, M., Reiners, P., Guanter, L., Segl, K., Chabrillat, S., Scheffler, D., Soppa, M.A., Bohn, N., Gorrono, J., Kokhanovsky, A., Bracher, A., Bachmann, M., Pato, M., Schneider, M., de los Reyes, R., Langheinrich, M., Holzwarth, S., Carmona, E., Storch, T., Pinnel, N., Habermeyer, M., Kokaly, R., Ong, C., Green, R.O., Moreno, J., Gascon, F., Hank, T., Ben-Dor, E., Di Mauro, B., Colombo, R., Milewski, R., Brede, B., Schmid, T., Anderson, C., Schickling, A., Krieger, V., Bock, M., La Porta, L., Fischer, S., 2026. Assessment of EnMAP data quality through global product validation activities. Remote Sensing of Environment 342, 115459. 10.1016/j.rse.2026.115459

Cavender-Bares, J., Schneider, F.D., Santos, M.J., Armstrong, A., Carnaval, A., Dahlin, K.M., Fatoyinbo, L., Hurtt, G.C., Schimel, D., Townsend, P.A., Ustin, S.L., Wang, Z., Wilson, A.M., 2022. Integrating remote sensing with ecology and evolution to advance biodiversity conservation. Nature Ecology & Evolution 6, 506–519. 10.1038/s41559-022-01702-5

Cimoli, E., Lucieer, A., Malenovský, Z., Woodgate, W., Janoutová, R., Turner, D., Haynes, R.S., Phinn, S., 2024. Mapping functional diversity of canopy physiological traits using UAS imaging spectroscopy. Remote Sensing of Environment 302, 113958. 10.1016/j.rse.2023.113958

Cogliati, S., Sarti, F., Chiarantini, L., Cosi, M., Lorusso, R., Lopinto, E., Miglietta, F., Genesio, L., Guanter, L., Damm, A., Pérez-López, S., Scheffler, D., Tagliabue, G., Panigada, C., Rascher, U., Dowling, T.P.F., Giardino, C., Colombo, R., 2021. The PRISMA imaging spectroscopy mission: overview and first performance analysis. Remote Sensing of Environment 262, 112499. 10.1016/j.rse.2021.112499

Corner, B.R., Mohan Narayanan, R., Reichenbach, S.E., 2000. Noise reduction in remote sensing imagery using data masking and principal component analysis. Presented at the Proc.SPIE, pp. 1–11. 10.1117/12.411533

de Bello, F., Lavergne, S., Meynard, C.N., Lepš, J., Thuiller, W., 2010. The partitioning of diversity: showing Theseus a way out of the labyrinth. Journal of Vegetation Science 21, 992–1000. 10.1111/j.1654-1103.2010.01195.x

de los Reyes, R., Langheinrich, M., Schwind, P., Richter, R., Pflug, B., Bachmann, M., Müller, R., Carmona, E., Zekoll, V., Reinartz, P., 2020. PACO: Python-Based Atmospheric Correction. Sensors 20, 1428. 10.3390/s20051428

de Sá, N.C., Baratchi, M., Hauser, L.T., van Bodegom, P., 2021. Exploring the Impact of Noise on Hybrid Inversion of PROSAIL RTM on Sentinel-2 Data. Remote Sensing 13, 648. 10.3390/rs13040648

Díaz, S., Lavorel, S., Chapin, F.S., Tecco, P.A., Gurvich, D.E., Grigulis, K., 2007. Functional Diversity — at the Crossroads between Ecosystem Functioning and Environmental Filters, in: Canadell, J.G., Pataki, D.E., Pitelka, L.F. (Eds.), Terrestrial Ecosystems in a Changing World. Springer Berlin Heidelberg, Berlin, Heidelberg, pp. 81–91. 10.1007/978-3-540-32730-1_7

Dietrich, C.R., Newsam, G.N., 1997. Fast and Exact Simulation of Stationary Gaussian Processes through Circulant Embedding of the Covariance Matrix. SIAM J. Sci. Comput. 18, 1088– 1107. 10.1137/S1064827592240555

Feilhauer, H., Somers, B., van der Linden, S., 2017. Optical trait indicators for remote sensing of plant species composition: Predictive power and seasonal variability. Ecological Indicators 73, 825–833. 10.1016/j.ecolind.2016.11.003

Féret, J.B., le Maire, G., Jay, S., Berveiller, D., Bendoula, R., Hmimina, G., Cheraiet, A., Oliveira, J.C., Ponzoni, F.J., Solanki, T., de Boissieu, F., Chave, J., Nouvellon, Y., Porcar-Castell, A., Proisy, C., Soudani, K., Gastellu-Etchegorry, J.P., Lefèvre-Fonollosa, M.J., 2019. Estimating leaf mass per area and equivalent water thickness based on leaf optical properties: Potential and limitations of physical modeling and machine learning. Remote Sensing of Environment 231, 110959. 10.1016/j.rse.2018.11.002

Forkel, M., Carvalhais, N., Schaphoff, S., v. Bloh, W., Migliavacca, M., Thurner, M., Thonicke, K., 2014. Identifying environmental controls on vegetation greenness phenology through model–data integration. Biogeosciences 11, 7025–7050. 10.5194/bg-11-7025-2014

Gamon, J.A., Somers, B., Malenovský, Z., Middleton, E.M., Rascher, U., Schaepman, M.E., 2019. Assessing Vegetation Function with Imaging Spectroscopy. Surveys in Geophysics. 10.1007/s10712-019-09511-5

Gastellu-Etchegorry, J.P., Martin, E., Gascon, F., 2004. DART: a 3D model for simulating satellite images and studying surface radiation budget. International Journal of Remote Sensing 25, 73–96. 10.1080/0143116031000115166

Gholizadeh, H., Gamon, J.A., Zygielbaum, A.I., Wang, R., Schweiger, A.K., Cavender-Bares, J., 2018. Remote sensing of biodiversity: Soil correction and data dimension reduction methods improve assessment of α-diversity (species richness) in prairie ecosystems. Remote Sensing of Environment 206, 240–253. 10.1016/j.rse.2017.12.014

Gómez-Dans, J.L., Lewis, P.E., Disney, M., 2016. Efficient Emulation of Radiative Transfer Codes Using Gaussian Processes and Application to Land Surface Parameter Inferences. Remote Sensing 8, 119. 10.3390/rs8020119

Gorroño, J., Guanter, L., Graf, L.V., Gascon, F., 2024. A Framework for the Estimation of Uncertainties and Spectral Error Correlation in Sentinel-2 Level-2A Data Products. IEEE Transactions on Geoscience and Remote Sensing 62, 1–13. 10.1109/TGRS.2024.3435021

Graf, L.V., Gorroño, J., Hueni, A., Walter, A., Aasen, H., 2023. Propagating Sentinel-2 Top-of-Atmosphere Radiometric Uncertainty Into Land Surface Phenology Metrics Using a Monte Carlo Framework. IEEE Journal of Selected Topics in Applied Earth Observations and Remote Sensing 16, 8632–8654. 10.1109/JSTARS.2023.3297713

Hauser, L.T., Féret, J.-B., An Binh, N., van der Windt, N., Sil, Â.F., Timmermans, J., Soudzilovskaia, N.A., van Bodegom, P.M., 2021a. Towards scalable estimation of plant functional diversity from Sentinel-2: In-situ validation in a heterogeneous (semi-)natural landscape. Remote Sensing of Environment 262, 112505. 10.1016/j.rse.2021.112505

Hauser, L.T., Timmermans, J., van der Windt, N., Sil, Â.F., César de Sá, N., Soudzilovskaia, N.A., van Bodegom, P.M., 2021b. Explaining discrepancies between spectral and in-situ plant diversity in multispectral satellite earth observation. Remote Sensing of Environment 265, 112684. 10.1016/j.rse.2021.112684

Helfenstein, I.S., Schneider, F.D., Schaepman, M.E., Morsdorf, F., 2022. Assessing biodiversity from space: Impact of spatial and spectral resolution on trait-based functional diversity. Remote Sensing of Environment 275, 113024. 10.1016/j.rse.2022.113024

Henniger, H., Bohn, F.J., Schmidt, K., Huth, A., 2023. A New Approach Combining a Multilayer Radiative Transfer Model with an Individual-Based Forest Model: Application to Boreal Forests in Finland. Remote Sensing 15, 3078. 10.3390/rs15123078

Hueni, A., Damm, A., Kneubuehler, M., Schläpfer, D., Schaepman, M.E., 2017. Field and Airborne Spectroscopy Cross Validation—Some Considerations. IEEE Journal of Selected Topics in Applied Earth Observations and Remote Sensing 10, 1117–1135. 10.1109/JSTARS.2016.2593984

Jacquemoud, S., Baret, F., 1990. PROSPECT: A model of leaf optical properties spectra. Remote Sensing of Environment 34, 75–91. 10.1016/0034-4257(90)90100-Z

JCGM, 2008. Evaluation of measurement data — Guide to the expression of uncertainty in measurement.

K. Thome, 2016. Calibration/validation error budgets, uncertainties, traceability and their importance to imaging spectrometry, in: 2016 IEEE International Geoscience and Remote Sensing Symposium (IGARSS). Presented at the 2016 IEEE International Geoscience and Remote Sensing Symposium (IGARSS), pp. 1912–1915. 10.1109/IGARSS.2016.7729492

Karasiak, N., Dejoux, J.-F., Fauvel, M., Willm, J., Monteil, C., Sheeren, D., 2019. Statistical Stability and Spatial Instability in Mapping Forest Tree Species by Comparing 9 Years of Satellite Image Time Series. Remote Sensing 11, 2512. 10.3390/rs11212512

Khintchine, A., 1934. Korrelationstheorie der stationären stochastischen Prozesse. Mathematische Annalen 109, 604–615. 10.1007/BF01449156

Kriegler, F.J., Malila, W.A., Nalepka, R.F., Richardson, W., 1969. Preprocessing Transformations and Their Effects on Multispectral Recognition, in: Remote Sensing of Environment, VI. p. 97.

Laliberté, E., Legendre, P., 2010. A distance-based framework for measuring functional diversity from multiple traits. Ecology 91, 299–305. 10.1890/08-2244.1

Li, Z., Leighton, H.G., 1992. Narrowband to Broadband Conversion with Spatially Autocorrelated Reflectance Measurements. Journal of Applied Meteorology and Climatology 31, 421–432. 10.1175/1520-0450(1992)031%3C0421:NTBCWS%3E2.0.CO;2

Ludwig, A., Doktor, D., Feilhauer, H., 2024. Is spectral pixel-to-pixel variation a reliable indicator of grassland biodiversity? A systematic assessment of the spectral variation hypothesis using spatial simulation experiments. Remote Sensing of Environment 302, 113988. 10.1016/j.rse.2023.113988

Ma, X., Migliavacca, M., Wirth, C., Bohn, J.F., Huth, A., Richter, R., Mahecha, D.M., 2020. Monitoring Plant Functional Diversity Using the Reflectance and Echo from Space. Remote Sensing 12. 10.3390/rs12081248

Magneville, C., Loiseau, N., Albouy, C., Casajus, N., Claverie, T., Escalas, A., Leprieur, F., Maire, E., Mouillot, D., Villéger, S., 2022. mFD: an R package to compute and illustrate the multiple facets of functional diversity. Ecography 2022. 10.1111/ecog.05904

Májeková, M., Paal, T., Plowman, N.S., Bryndová, M., Kasari, L., Norberg, A., Weiss, M., Bishop, T.R., Luke, S.H., Sam, K., Le Bagousse-Pinguet, Y., Lepš, J., Götzenberger, L., de Bello, F., 2016. Evaluating Functional Diversity. PLoS One 11. 10.1371/journal.pone.0149270

Mannschatz, T., Pflug, B., Borg, E., Feger, K.-H., Dietrich, P., 2014. Uncertainties of LAI estimation from satellite imaging due to atmospheric correction. Remote Sensing of Environment 153, 24–39. 10.1016/j.rse.2014.07.020

Martínez-Ferrer, L., Moreno-Martínez, Á., Campos-Taberner, M., García-Haro, F.J., Muñoz-Marí, J., Running, S.W., Kimball, J., Clinton, N., Camps-Valls, G., 2022. Quantifying uncertainty in high resolution biophysical variable retrieval with machine learning. Remote Sensing of Environment 280, 113199. 10.1016/j.rse.2022.113199

Norbert Wiener, 1930. Generalized harmonic analysis. Acta Mathematica 55, 117–258. 10.1007/BF02546511

Ollinger, S.V., 2011. Sources of variability in canopy reflectance and the convergent properties of plants. New Phytologist 189, 375–394. 10.1111/j.1469-8137.2010.03536.x

Pacheco-Labrador, J., 2026. Code for: Assessing the impact of data uncertainty on remote sensing estimation of plant functional diversity. 10.5281/zenodo.22872340

Pacheco-Labrador, J., de Bello, F., Migliavacca, M., Ma, X., Carvalhais, N., Wirth, C., 2023. A generalizable normalization for assessing plant functional diversity metrics across scales from remote sensing. Methods in Ecology and Evolution 14, 2123–2136. 10.1111/2041-210X.14163

Pacheco-Labrador, J., Gomarasca, U., Li, W., Migliavacca, M., Jung, M., Duveiller, G., 2025. BOSSE v1.0: the Biodiversity Observing System Simulation Experiment. Geosci. Model Dev. 18, 8401–8422. 10.5194/gmd-18-8401-2025

Pacheco-Labrador, J., Gomarasca, U., Pabon-Moreno, D.E., Li, W., Migliavacca, M., Jung, M., Duveiller, G., 2026. Benchmarking remote sensing methods to capture plant functional diversity from space. Ecological Informatics 94, 103636. 10.1016/j.ecoinf.2026.103636

Pacheco-Labrador, J., Gomarasca, U., Webber, U., Li, W., Hamdi, Z., Pabon, D., Loos, D., Jung, M., Duveiller, G., 2024. BOSSE: The Biodiversity Observing System Simulation Experiment for Remote Sensing. Geophysical research abstracts. 10.5194/egusphere-egu24-6746

Pacheco-Labrador, J., Migliavacca, M., Ma, X., Mahecha, M.D., Carvalhais, N., Weber, U., Benavides, R., Bouriaud, O., Barnoaiea, I., Coomes, D.A., Bohn, F.J., Kraemer, G., Heiden, U., Huth, A., Wirth, C., 2022. Challenging the link between functional and spectral diversity with radiative transfer modeling and data. Remote Sensing of Environment 280, 113170. 10.1016/j.rse.2022.113170

Pacheco-Labrador, J., Weber, U., Li, W., Gomarasca, U., Pabon-Moreno, D.E., Migliavacca, M., Martin, J., Duveiller, G., 2025. The Biodiversity Observing System Simulation Experiment (BOSSE v1.0) ERA5-Land Meteorological time series. 10.5281/zenodo.14717038

Perrone, M., Conti, L., Galland, T., Komárek, J., Lagner, O., Torresani, M., Rossi, C., Carmona, C.P., de Bello, F., Rocchini, D., Moudrý, V., Šímová, P., Bagella, S., Malavasi, M., 2024. “Flower power”: How flowering affects spectral diversity metrics and their relationship with plant diversity. Ecological Informatics 81, 102589. 10.1016/j.ecoinf.2024.102589

Raiho, A.M., Cawse-Nicholson, K., Chlus, A., Dozier, J., Gierach, M., Miner, K., Schneider, F., Schimel, D., Serbin, S., Shiklomanov, A.N., Thompson, D.R., Townsend, P.A., Zareh, S., Skiles, M., Poulter, B., 2023. Exploring Mission Design for Imaging Spectroscopy Retrievals for Land and Aquatic Ecosystems. Journal of Geophysical Research: Biogeosciences 128, e2022JG006833. 10.1029/2022JG006833

Ricotta, C., Szeidl, L., 2006. Towards a unifying approach to diversity measures: Bridging the gap between the Shannon entropy and Rao’s quadratic index. Theoretical Population Biology 70, 237–243. 10.1016/j.tpb.2006.06.003

Rossi, C., Gholizadeh, H., 2023. Uncovering the hidden: Leveraging sub-pixel spectral diversity to estimate plant diversity from space. Remote Sensing of Environment 296, 113734. 10.1016/j.rse.2023.113734

Rossi, C., Hueni, A., Koch, T.L., Pacheco-Labrador, J., Karaman, K., Abdullah, H.J., Darvishzadeh, R., Gholizadeh, H., Féret, J.-B., Torresani, M., Vihervaara, P., Santos, M.J., in press. Uncertainties in optical remote sensing of plant diversity. Remote Sensing of Environment. 10.1016/j.rse.2026.115688

Rossi, C., Kneubühler, M., Schütz, M., Schaepman, M.E., Haller, R.M., Risch, A.C., 2021. Spatial resolution, spectral metrics and biomass are key aspects in estimating plant species richness from spectral diversity in species-rich grasslands. Remote Sensing in Ecology and Conservation n/a. 10.1002/rse2.244

Rossi, C., McMillan, N.A., Schweizer, J.M., Gholizadeh, H., Groen, M., Ioannidis, N., Hauser, L.T., 2024. Parcel level temporal variance of remotely sensed spectral reflectance predicts plant diversity. Environmental Research Letters 19, 074023. 10.1088/1748-9326/ad545a

Schneider, F.D., Morsdorf, F., Schmid, B., Petchey, O.L., Hueni, A., Schimel, D.S., Schaepman, M.E., 2017. Mapping functional diversity from remotely sensed morphological and physiological forest traits. Nature Communications 8, 1441. 10.1038/s41467-017-01530-3

Skidmore, A.K., Coops, N.C., Neinavaz, E., Ali, A., Schaepman, M.E., Paganini, M., Kissling, W.D., Vihervaara, P., Darvishzadeh, R., Feilhauer, H., Fernandez, M., Fernández, N., Gorelick, N., Geijzendorffer, I., Heiden, U., Heurich, M., Hobern, D., Holzwarth, S., Muller-Karger, F.E., Van De Kerchove, R., Lausch, A., Leitão, P.J., Lock, M.C., Mücher, C.A., O’Connor, B., Rocchini, D., Roeoesli, C., Turner, W., Vis, J.K., Wang, T., Wegmann, M., Wingate, V., 2021. Priority list of biodiversity metrics to observe from space. Nature Ecology & Evolution 5, 896–906. 10.1038/s41559-021-01451-x

Smith, G.M., Milton, E.J., 1999. The use of the empirical line method to calibrate remotely sensed data to reflectance. International Journal of Remote Sensing 20, 2653–2662. 10.1080/014311699211994

Storch, T., Honold, H.-P., Chabrillat, S., Habermeyer, M., Tucker, P., Brell, M., Ohndorf, A., Wirth, K., Betz, M., Kuchler, M., Mühle, H., Carmona, E., Baur, S., Mücke, M., Löw, S., Schulze, D., Zimmermann, S., Lenzen, C., Wiesner, S., Aida, S., Kahle, R., Willburger, P., Hartung, S., Dietrich, D., Plesia, N., Tegler, M., Schork, K., Alonso, K., Marshall, D., Gerasch, B., Schwind, P., Pato, M., Schneider, M., de los Reyes, R., Langheinrich, M., Wenzel, J., Bachmann, M., Holzwarth, S., Pinnel, N., Guanter, L., Segl, K., Scheffler, D., Foerster, S., Bohn, N., Bracher, A., Soppa, M.A., Gascon, F., Green, R., Kokaly, R., Moreno, J., Ong, C., Sornig, M., Wernitz, R., Bagschik, K., Reintsema, D., La Porta, L., Schickling, A., Fischer, S., 2023. The EnMAP imaging spectroscopy mission towards operations. Remote Sensing of Environment 294, 113632. 10.1016/j.rse.2023.113632

Torresani, M., Rossi, C., Perrone, M., Hauser, L.T., Féret, J.-B., Moudrý, V., Simova, P., Ricotta, C., Foody, G.M., Kacic, P., Feilhauer, H., Malavasi, M., Tognetti, R., Rocchini, D., 2024. Reviewing the spectral variation hypothesis: Twenty years in the tumultuous sea of biodiversity estimation by remote sensing. Ecological Informatics 102702. 10.1016/j.ecoinf.2024.102702

Ustin, S.L., Gitelson, A.A., Jacquemoud, S., Schaepman, M., Asner, G.P., Gamon, J.A., Zarco-Tejada, P., 2009. Retrieval of foliar information about plant pigment systems from high resolution spectroscopy. Remote Sensing of Environment 113, S67–S77. 10.1016/j.rse.2008.10.019

van der Tol, C., Verhoef, W., Timmermans, J., Verhoef, A., Su, Z., 2009. An integrated model of soil-canopy spectral radiances, photosynthesis, fluorescence, temperature and energy balance. Biogeosciences 6, 3109–3129. 10.5194/bg-6-3109-2009

Vasquez, V.R., Whiting, W.B., 2005. Accounting for Both Random Errors and Systematic Errors in Uncertainty Propagation Analysis of Computer Models Involving Experimental Measurements with Monte Carlo Methods. Risk Analysis 25, 1669–1681. 10.1111/j.1539-6924.2005.00704.x

Verhoef, W., 1984. Light scattering by leaf layers with application to canopy reflectance modeling: The SAIL model. Remote Sensing of Environment 16, 125–141. 10.1016/0034-4257(84)90057-9

Verrelst, J., Rivera, J.P., van der Tol, C., Magnani, F., Mohammed, G., Moreno, J., 2015. Global sensitivity analysis of the SCOPE model: What drives simulated canopy-leaving sun-induced fluorescence? Remote Sensing of Environment 166, 8–21. 10.1016/j.rse.2015.06.002

Wallis, C.I.B., Crofts, A.L., Jackisch, R., Kothari, S., Tougas, G., Arroyo-Mora, J.P., Hacker, P., Coops, N., Kalacska, M., Laliberté, E., Vellend, M., 2025. Methodological considerations for studying spectral-plant diversity relationships. Remote Sensing of Environment 328, 114907. 10.1016/j.rse.2025.114907

Wang, R., Gamon, J.A., 2019. Remote sensing of terrestrial plant biodiversity. Remote Sensing of Environment 231, 111218. 10.1016/j.rse.2019.111218

Wang, R., Gamon, J.A., Cavender-Bares, J., Townsend, P.A., Zygielbaum, A.I., 2018a. The spatial sensitivity of the spectral diversity–biodiversity relationship: an experimental test in a prairie grassland. Ecological Applications 28, 541–556. 10.1002/eap.1669

Wang, R., Gamon, J.A., Schweiger, A.K., Cavender-Bares, J., Townsend, P.A., Zygielbaum, A.I., Kothari, S., 2018b. Influence of species richness, evenness, and composition on optical diversity: A simulation study. Remote Sensing of Environment 211, 218–228. 10.1016/j.rse.2018.04.010

Wang, Z., Townsend, P.A., Schweiger, A.K., Couture, J.J., Singh, A., Hobbie, S.E., Cavender-Bares, J., 2019a. Mapping foliar functional traits and their uncertainties across three years in a grassland experiment. Remote Sensing of Environment 221, 405–416. 10.1016/j.rse.2018.11.016

Wang, Z., Townsend, P.A., Schweiger, A.K., Couture, J.J., Singh, A., Hobbie, S.E., Cavender-Bares, J., 2019b. Mapping foliar functional traits and their uncertainties across three years in a grassland experiment. Remote Sensing of Environment 221, 405–416. 10.1016/j.rse.2018.11.016

Widlowski, J.-L., 2015. Conformity testing of satellite-derived quantitative surface variables. Environmental Science & Policy 51, 149–169. 10.1016/j.envsci.2015.03.018

Wood, A.T.A., Chan, G., 1994. Simulation of Stationary Gaussian Processes in [0, 1] d. Journal of Computational and Graphical Statistics 3, 409–432. 10.1080/10618600.1994.10474655

Zhang, M., Ibrahim, A., Franz, B.A., Sayer, A.M., Werdell, P.J., McKinna, L.I., 2024. Spectral correlation in MODIS water-leaving reflectance retrieval uncertainty. Opt. Express 32, 2490–2506. 10.1364/OE.502561

Zhang, Y., Migliavacca, M., Penuelas, J., Ju, W., 2021. Advances in hyperspectral remote sensing of vegetation traits and functions. Remote Sensing of Environment 252, 112121. 10.1016/j.rse.2020.112121

Zhu, L., Fu, B., Zhu, H., Wang, C., Jiao, L., Zhou, J., 2017. Trait choice profoundly affected the ecological conclusions drawn from functional diversity measures. Scientific Reports 7, 3643. 10.1038/s41598-017-03812-8

Zhu, Y., Wang, Q.A., Li, W., Cai, X., 2018. Analytic uncertainty and sensitivity analysis of models with input correlations. Physica A: Statistical Mechanics and its Applications 494, 140–162. 10.1016/j.physa.2017.12.041

