## Supplementary Material for "Assessing the impact of data uncertainty on remote sensing estimation of plant functional diversity"

### Supplementary Material for:
### Assessing the impact of data uncertainty on remote sensing estimation of plant functional diversity

Javier Pacheco Labrador[1], Christian Rossi[2,3], and Maria J. Santos[2]

[1] Environmental Remote Sensing and Spectroscopy Laboratory (SpecLab), Institute of Agricultural Sciences (ICA), Spanish National Research Council (CSIC), Madrid, Spain

[2] Department of Geography, University of Zurich, Winterthurerstrasse 190, 8057 Zürich, Switzerland

[3] Department of Geoinformation, Swiss National Park, Runatsch 124-Chastè Planta-Wildenberg, 7530 Zernez, Switzerland

**Figure S1**

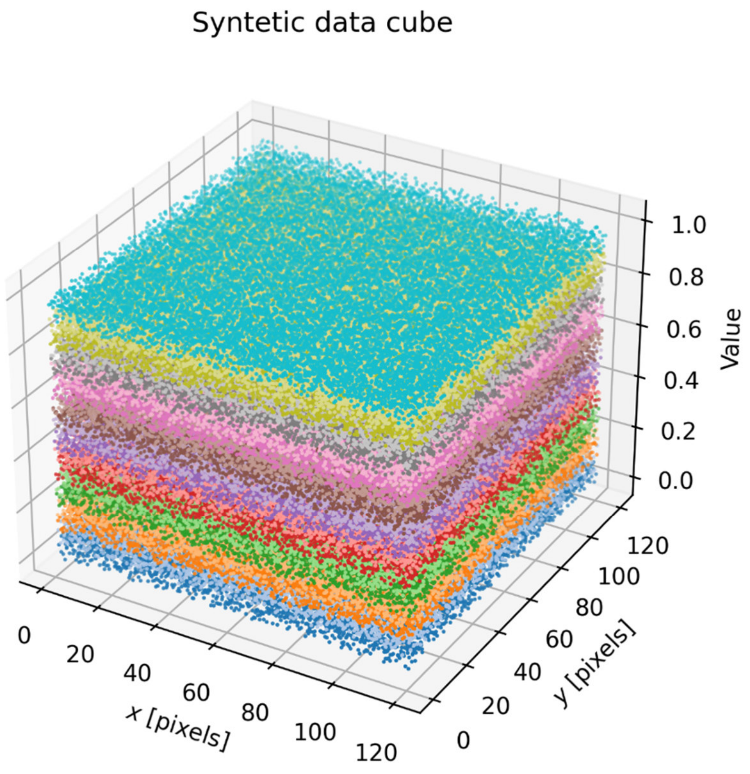


Figure S1. Synthetic data cube of random values organized by layers used to exemplify the application of uncertainties.

**Figure S2**

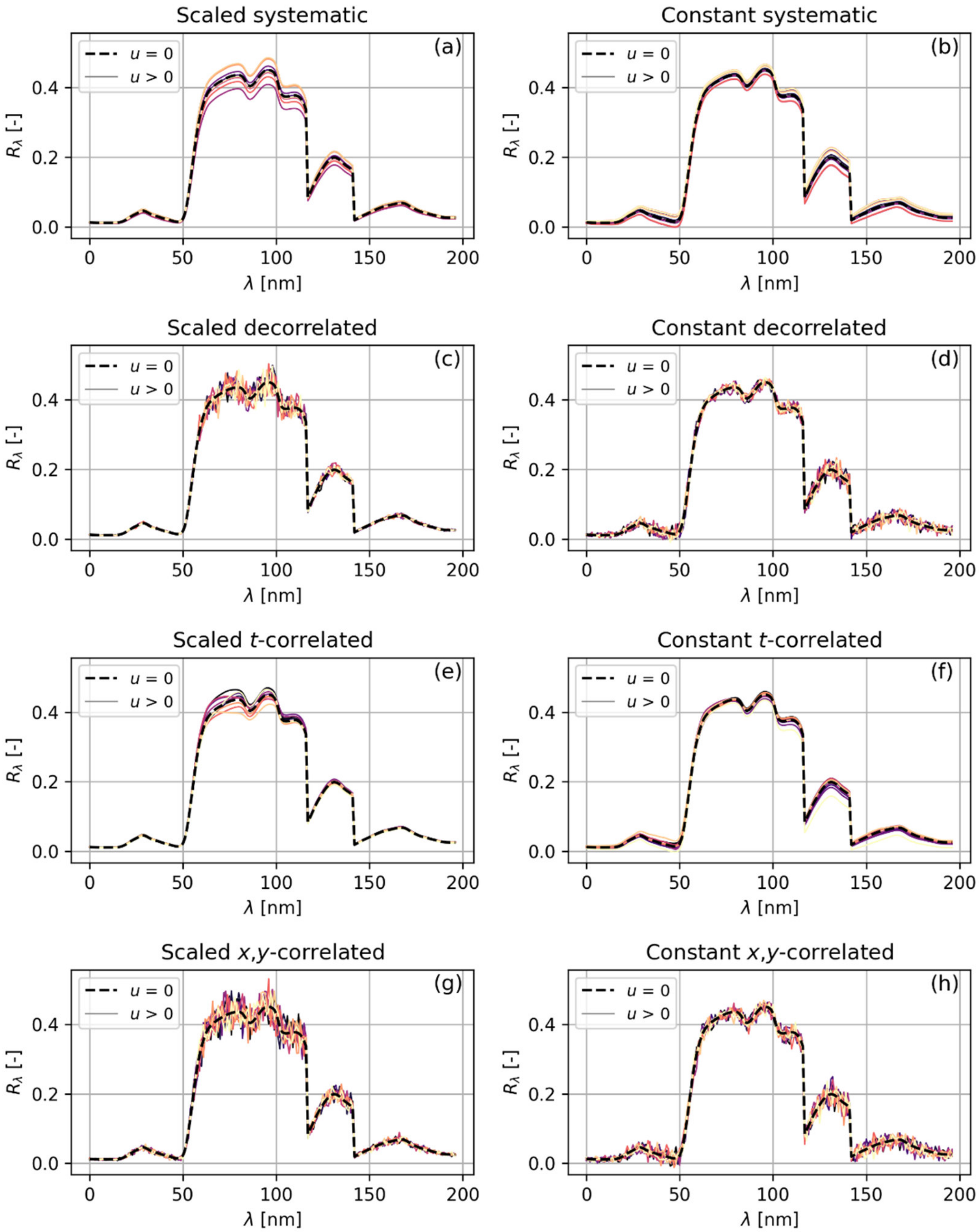


Figure S2. Example of spectral reflectance uncertainty types simulated from five realizations.

**Figure S3**

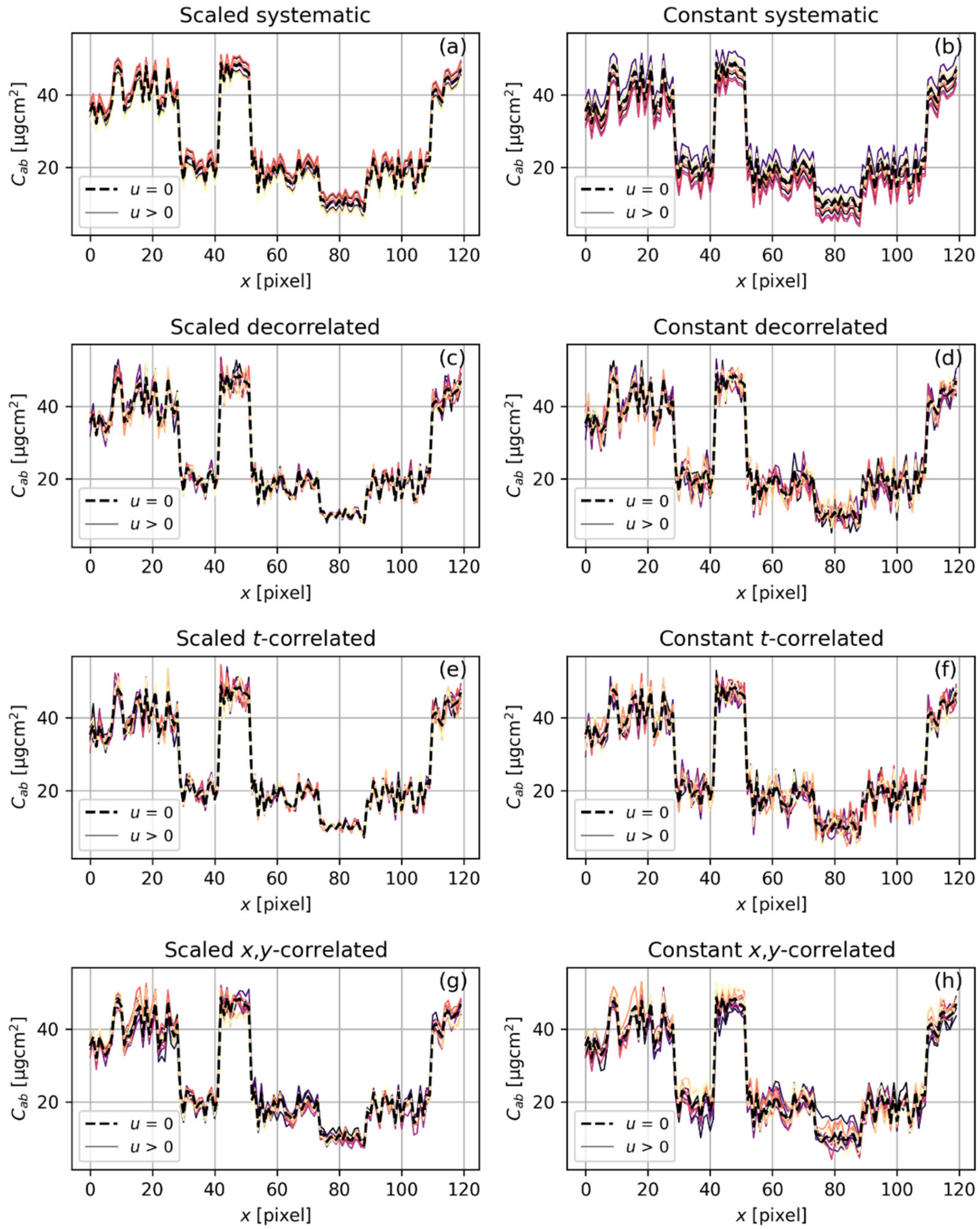


Figure S3. Example of leaf chlorophyll content uncertainty types across a trait map row simulated from five realizations. Leaf chlorophyll content values are presented along a row of the image.

**Figure S4**

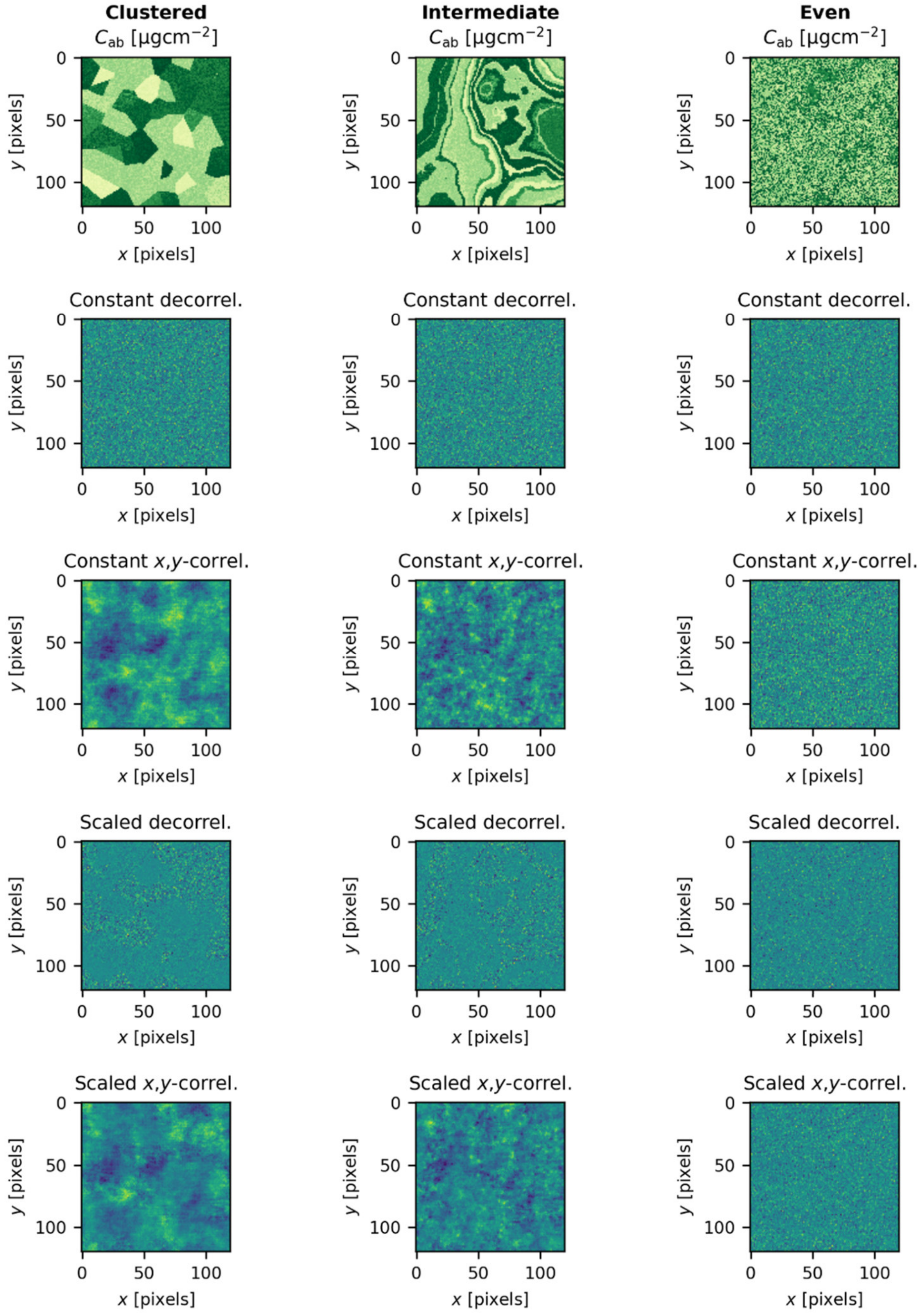


Figure S4. Example of leaf chlorophyll content and the random decorrelated and spatially-correlated uncertainties.

### Figure S5

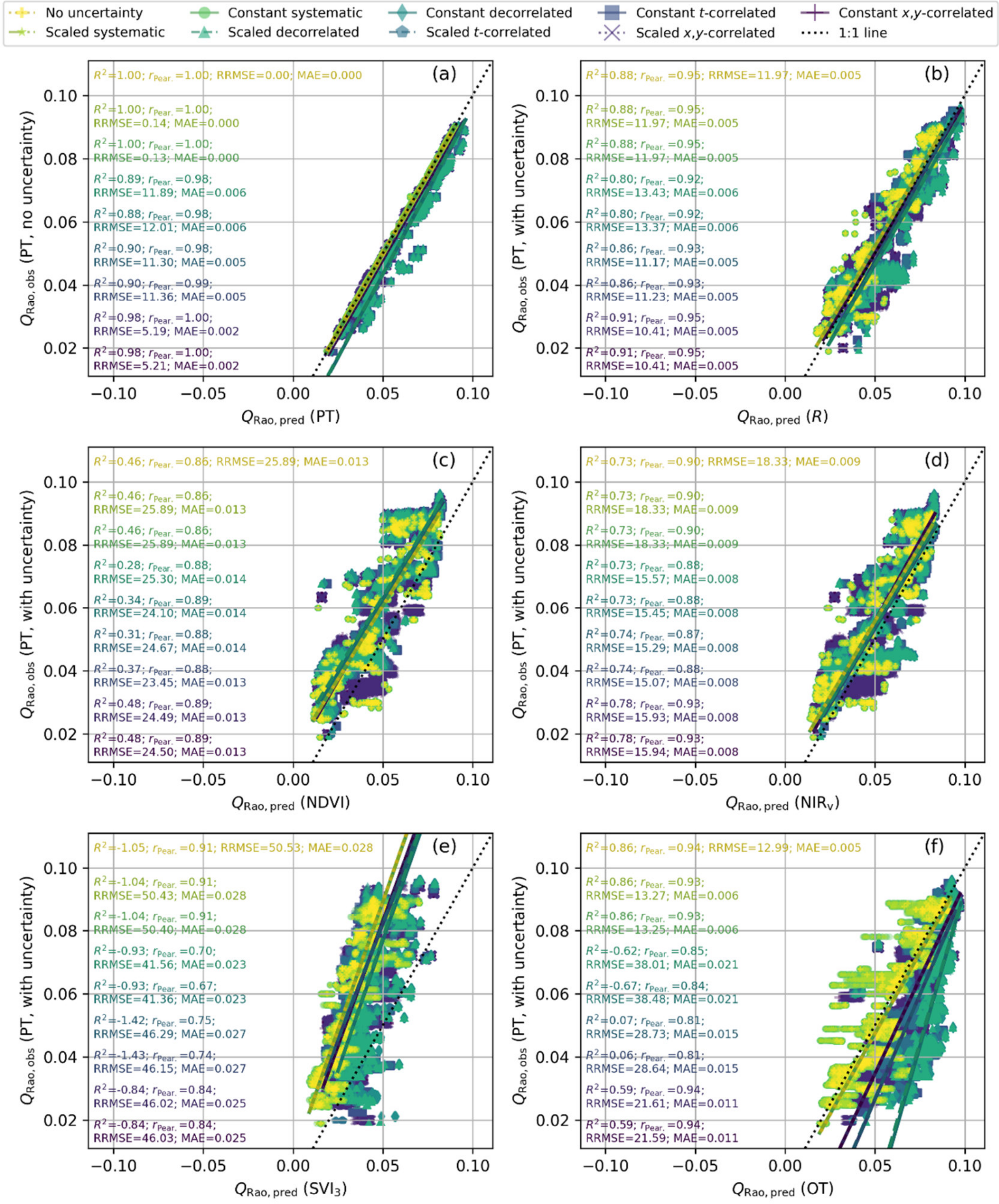


Figure S5. Comparison of Rao's quadratic entropy ($Q_{\mathrm{Rao}}$) computed from plant traits without uncertainty and from plant traits (a), reflectance factors (b), normalized difference vegetation index (c), near infrared reflectance of vegetation (d), a cube of three spectral indices (e), and optical traits (f) featuring different data uncertainty types ("apparent error").

#### Figure S6

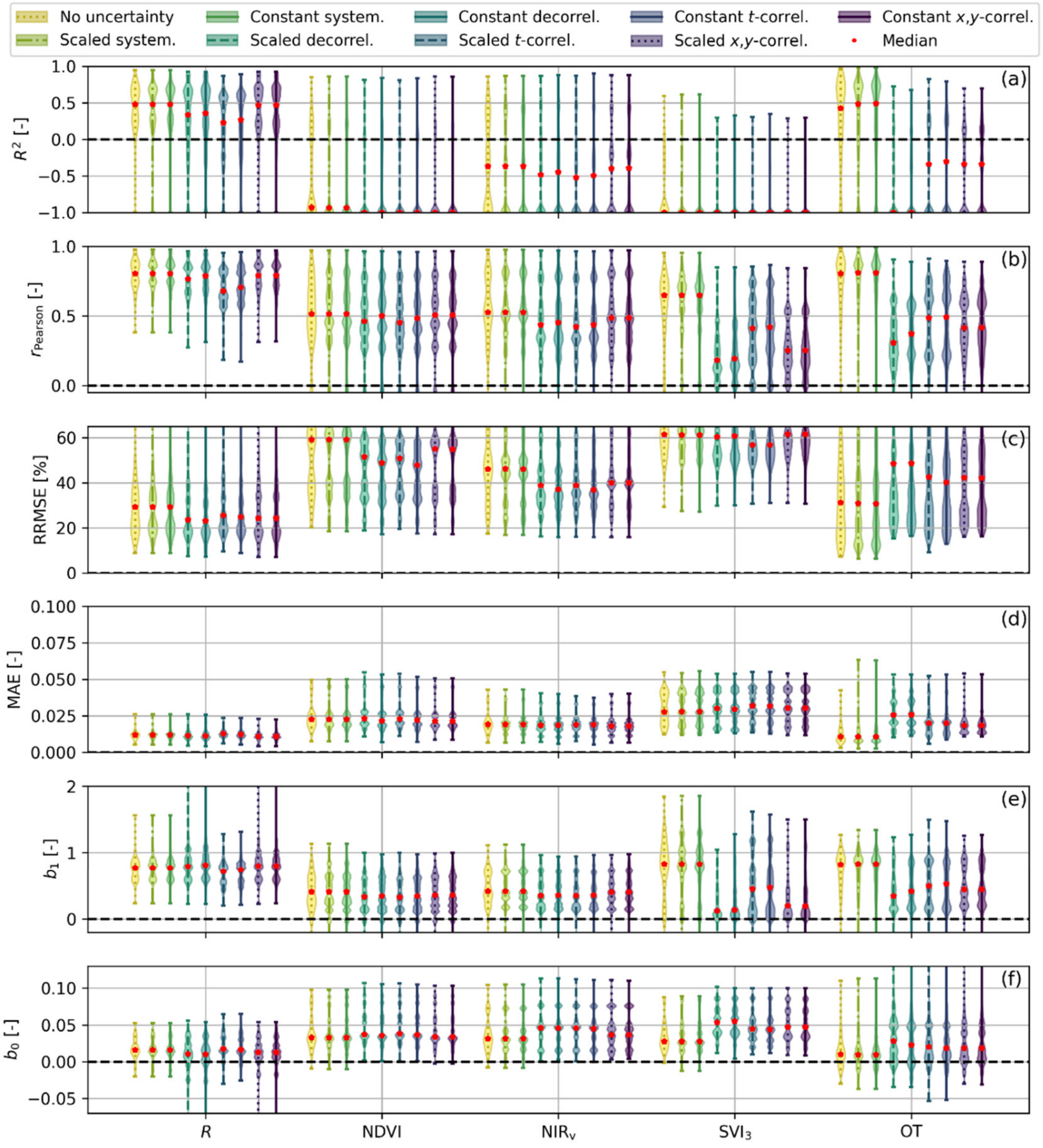


Figure S6. "Apparent error" statistics of the performance in the estimation of plant functional diversity for the different types of uncertainty: (a) coefficient of determination, (b) Pearson correlation coefficient, (c) relative root mean squared error, (d) mean absolute error, slope (e) and offset (f) of the predicted-observed relationship.

**Figure S7**

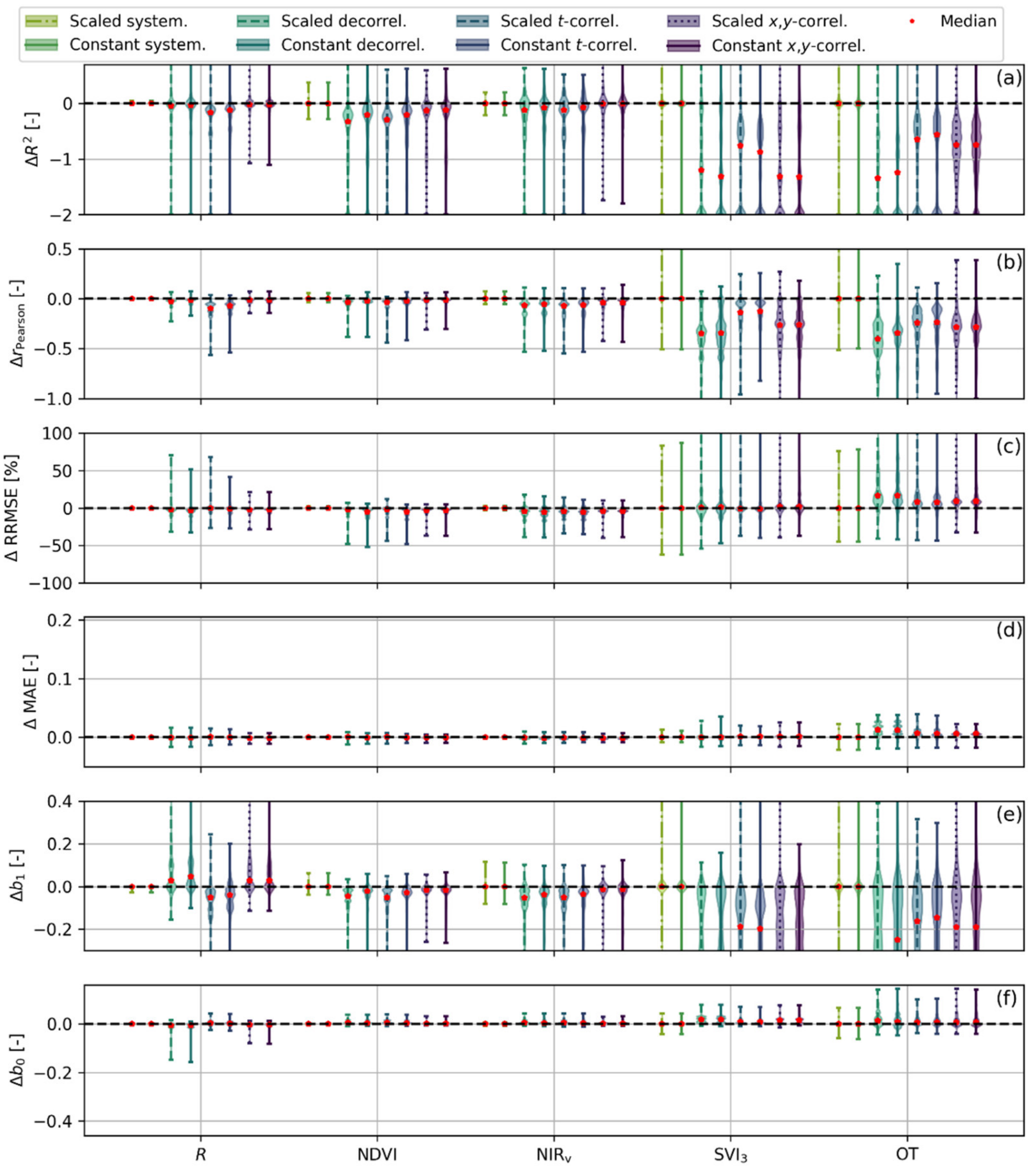


Figure S7. Difference of "apparent error" statistics of the performance in the estimation of plant functional diversity with the case "no uncertainty" for the different types of uncertainty: (a) coefficient of determination, (b) Pearson correlation coefficient, (c) relative root mean squared error, (d) mean absolute error, slope (e), and offset (f) of the predicted-observed relationship.

#### Figure S8

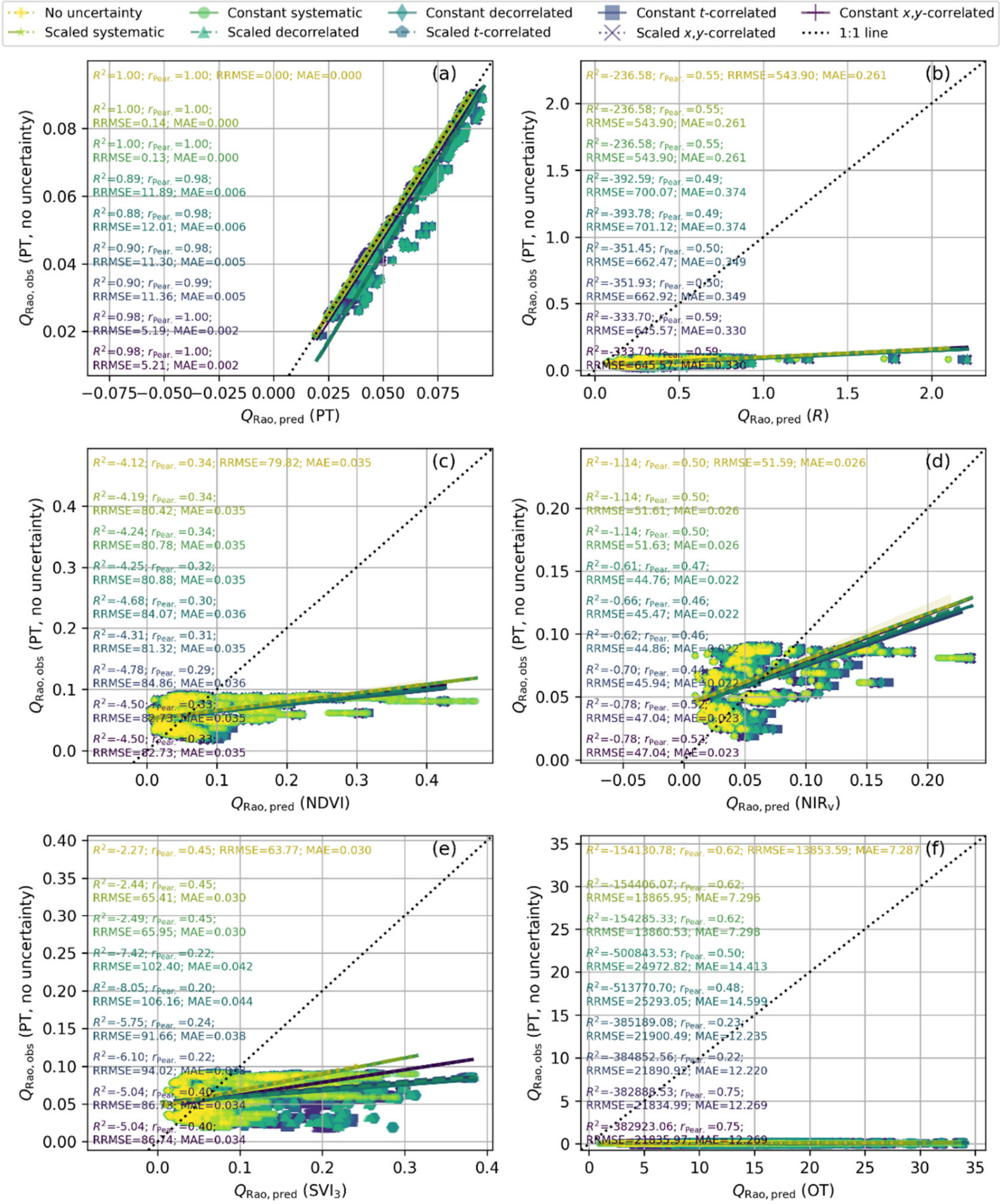


Figure S8. Comparison of Rao's quadratic entropy ($Q_{Rao}$) computed from plant traits without uncertainty and from plant traits (a), reflectance factors (b), normalized difference vegetation index (c), near infrared reflectance of vegetation (d), a cube of three spectral indices (e), and optical traits (f) featuring different data uncertainty types ("true error"). No preprocessing (standardization and principal component analysis) was applied to traits before computing $Q_{Rao}$.

**Figure S9**

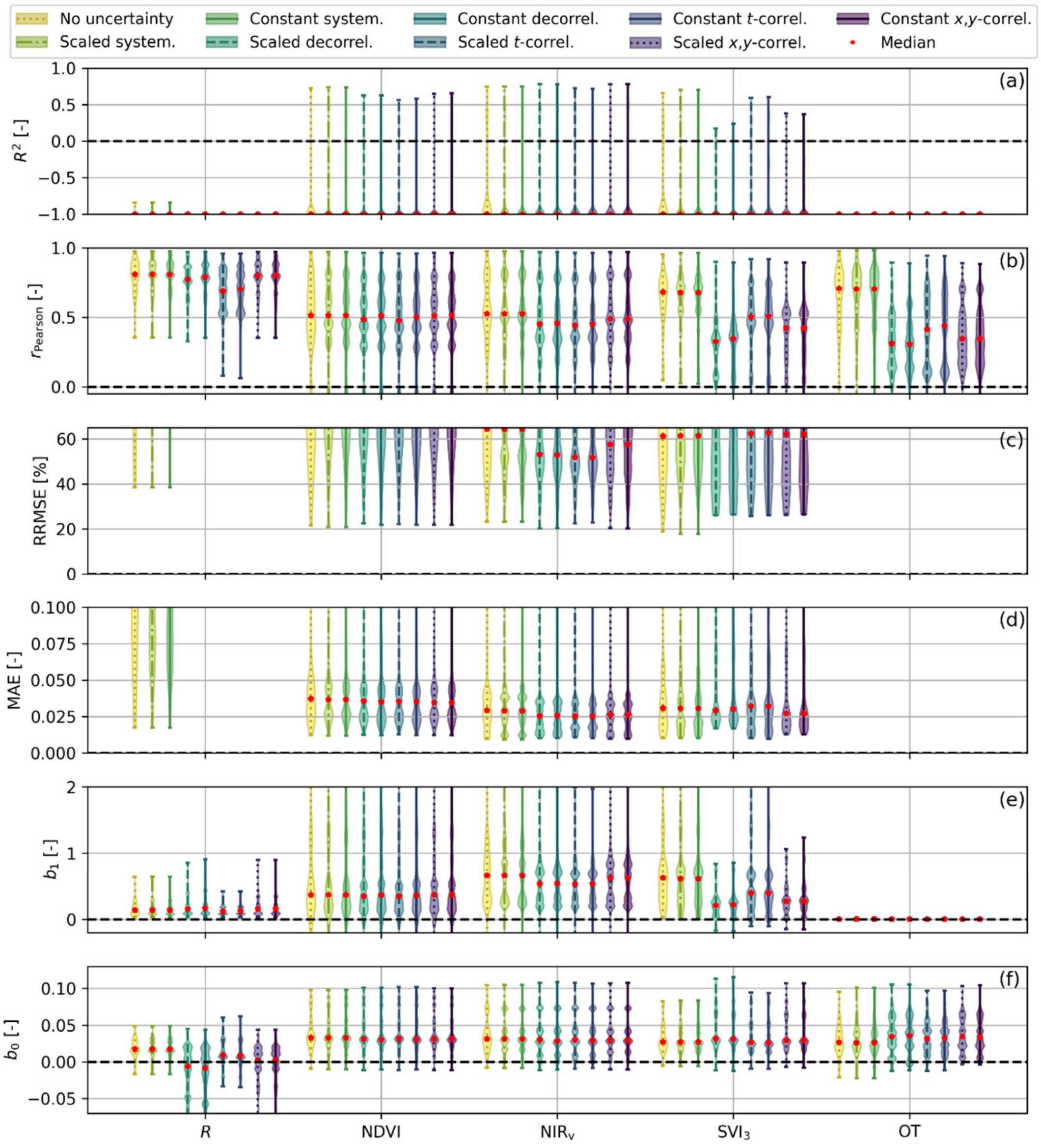


Figure S9. "True error" statistics of the performance in the estimation of plant functional diversity for the different types of uncertainty: (a) coefficient of determination, (b) Pearson correlation coefficient, (c) relative root mean squared error, (d) mean absolute error, slope (e) and offset (f) of the predicted-observed relationship. No preprocessing (standardization and principal component analysis) was applied to traits before computing $Q_{\text{Rao}}$.

**Figure S10**

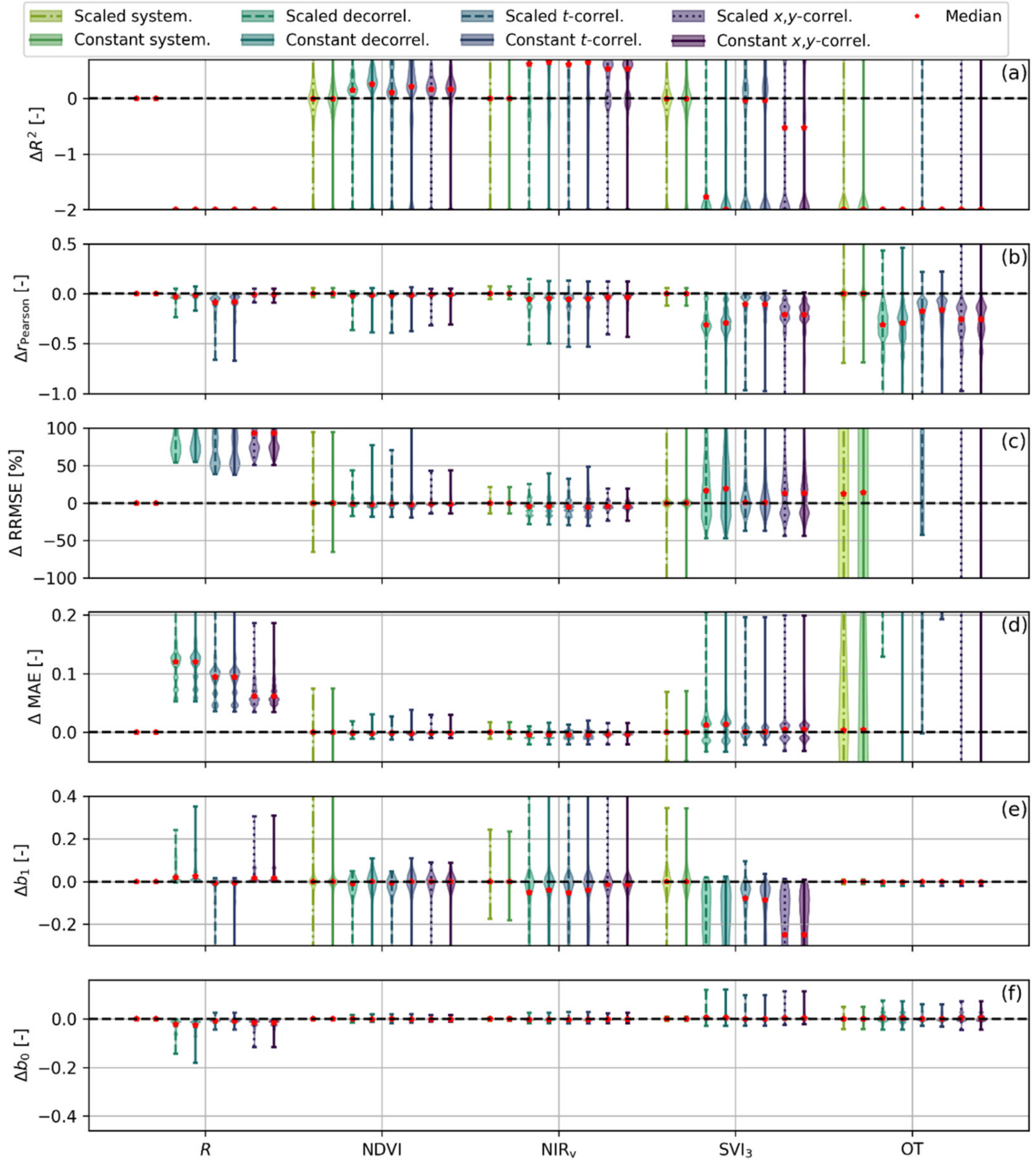


Figure S10. Difference of "true error" statistics of the performance in the estimation of plant functional diversity with the case "no uncertainty" for the different types of uncertainty: (a) coefficient of determination, (b) Pearson correlation coefficient, (c) relative root mean squared error, (d) mean absolute error, slope (e), and offset (f) of the predicted-observed relationship. No preprocessing (standardization and principal component analysis) was applied to traits before computing $Q_{\mathrm{Rao}}$.

**Figure S11**

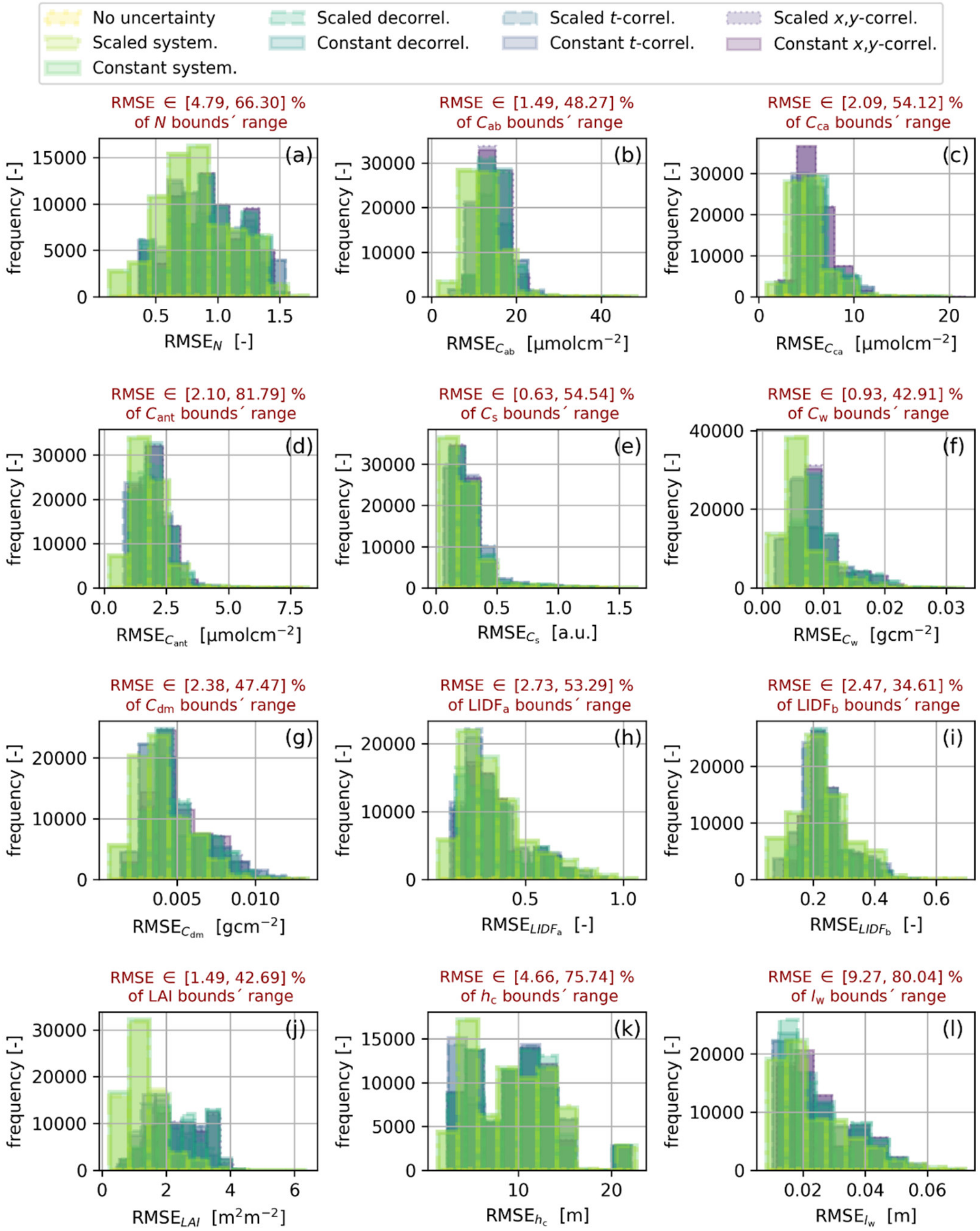


Figure S11. Optical trait retrieval root mean squared error (RMSE) distributions presented by trait and uncertainty type.
